# A Bayesian multidimensional approach to decipher the genetic basis of dynamic phenotypes in multiple species

**DOI:** 10.64898/2026.04.01.715770

**Authors:** Louis Blois, Benjamin Heuclin, Anthony Bernard, Marie Denis, Elisabeth Dirlewanger, Marie Foulongne-Oriol, Philippe Marullo, Emilien Peltier, José Quero-Garciá, Elisa Marguerit, Jean-Marc Gion

## Abstract

Deciphering the genetic architecture of complex quantitative phenotypes remains challenging in quantitative genetics. These traits not only depend of multiple genetic factors but are also established over time and environments. Although quantitative genetics has investigated the genetic determinism of phenotypic plasticity in contrasted environmental conditions, the time related phenotypic plasticity has received less attention.

Here we proposed a multivariate Bayesian framework, the Bayesian Varying Coefficient Model, designed for analysing the genetic architecture of the time related phenotypic plasticity by a multilocus approach. We applied the BVCM to time series phenotypes measured at various time scales (daily, monthly, yearly) across a diverse set of biological species. We included in this study: yeast (*Saccharomyces cerevisiae*), fungi (*Fusarium graminearum*), eucalyptus (*Eucalyptus urophylla × E. grandis*), and sweet cherry tree (*Prunus avium*). The BVCM results were compared with those obtained with a known genome-wide association method carried out time by time.

For all species and traits, the BVCM was able to detect the major QTL identified by marker-trait association methods and revealed additional genetic regions of weak effect. It also increased the phenotypic variance explained for most of the phenotypes considered. It revealed dynamic QTLs with transitory, increasing or decreasing effects over time. By considering both the temporal and genetic multivariate structures in a single statistical model, we increased our understanding of the genetic architecture of complex traits notably by reducing the issue of missing heritability. More broadly, this work raises the foundation for extended applications in functional genomics, evolutionary ecology, and crop breeding programs, in which time-related phenotypic plasticity remains crucial for predicting and selecting key quantitative complex traits.

**Key message:** By capturing the genetic factors influencing the time related phenotypic plasticity, our approach contributes to a deeper understanding of the dynamic nature of genotype–phenotype relationships.

## 1. Introduction

Understanding the genetic basis of complex quantitative traits remains a challenge due to their oligo/multi-genic determinism (Goddard *et al*., 2016). In agronomy, most quantitative traits are measured at the exploitation age considered as representative of the genotype performances, or after a fixed number of days following planting or stress events. However, phenotypic traits such as growth, developmental, and phenological processes are established over time and environments. Analyzing these dynamics require time series data of each phenotype. In the context of quantitative genetics, the study of time series offers the opportunity to reveal the effect of various genetic regions in the establishment of phenotypes and increases the accuracy and power of quantitative trait loci (QTL) detection approaches (Moore *et al*., 2013; Sang *et al*., 2019; Miao *et al*., 2020). While many efforts have been deployed in quantitative genetics to consider phenotypic plasticity for spatial environmental variation as reviewed by Laitinen & Nikoloski (2019), the time related phenotypic plasticity remains marginally explored and statistical approaches struggle to consider the multidimensional aspect of such data sets. In most of marker traits association studies, QTL mapping or genome-wide association studies (GWAS), two approaches were used to consider time series. Firstly, analyses were carried out for each time step of time series for plant height in rice (Yan *et al*., 1998) and wheat (Wu *et al*., 2010), waterlogging tolerance at maize seedling stages (Osman *et al*., 2013), water status in eucalyptus (Bartholomé *et al*., 2020), or canopy coverage in soybean (Li *et al*., 2023). However, the time-by-time analysis did not reflect the dependency of the data measured on each genotype for multiple time points. Secondly, in order to overcome this limitation, the temporal aspect of phenotype variation was incorporated by converting the phenotype into function parameters (Ma *et al*., 2002; Wu *et al*., 2004) or PCA/FPCA (principal component analysis/functional principal component analysis) coordinates (Kwak *et al*., 2016; Muraya *et al*., 2017; Xu *et al*., 2018b,a; Miao *et al*., 2020; Li *et al*., 2023). However, the complexity and informational content of the time series were not fully exploited by these approaches.

Quantitative and association mapping traditionally relied on single-locus approaches, which enabled identifying major QTL associated with many mendelian mono/oligo genic traits such as diseases resistance (St Clair, 2010). However, the polygenic genetic architecture of quantitative traits involves numerous loci with a wide range of effects from few strong effects to putative huge number of small effects, as well as potential interactions among them (Holland, 2007; Mackay, 2009). These aspects, which contribute significantly to phenotypic variation, are difficult to statistically identify (Hill, 2010). Widely used single-locus methods often lack the power to identify loci with small effects or their interactions (epistatic effects) (Carlson *et al*., 2004). Multilocus approaches were developed to address these limitations by incorporating more than one genomic region within a single analysis conducted for a wide range of statistical models. For instance, composite interval mapping (CIM; Jansen & Stam, 1994; Zeng, 1994) and the Bayesian-information and Linkage-disequilibrium Iteratively Nested Keyway method (BLINK method; Huang *et al*., 2018) represented multilocus strategies with contrasted levels of “multi-dimensions”. CIM involves a series of single-locus scans in which detected QTLs are included as cofactors in subsequent analyses. This improved detection power and resolution for loci with small effects, although it did not fully use genome-wide information (Segura *et al*., 2012). In contrast, BLINK adopted a more comprehensive multilocus framework by iteratively selecting an optimal subset of markers as covariates and updating them until the model converged (Huang *et al*., 2018). While multilocus models differed in the level of multivariate structure they exhibited, they consistently outperformed single-locus approaches capturing the complexity of genetic architectures (Hoggart *et al*., 2008; Segura *et al*., 2012; Xu *et al*., 2018c; Kaler *et al*., 2020; Zhong *et al*., 2021; Adhikari *et al*., 2023; Osorio-Guarin *et al*., 2024). In these studies, the advantages of multilocus approaches were the detection of more genetic markers associated with already known genetic regions and the decrease of the false positive and false negative discovery rate in association mapping.

Currently, multilocus statistical approaches were based on “stepwise like” variable selection methods. These methods were broadly contested because of bias in parameter estimations and inconsistencies among model selection (Steyerberg *et al*., 2000; Whittingham *et al*., 2006). Alternatively, Steyerberg *et al*. (2000) promoted the use of penalized likelihood methods such as the ridge regression (Hoerl & Kennard, 1970) in a context of multicollinearity, or the Lasso regression (Tibshirani, 1996) when the number of predictors was large relatively to the number of observations. The Bayesian approach provides a natural framework for variable selection by placing shrinkage priors on the regression coefficients, thereby introducing regularization (Bickel *et al*., 2006). Among them, the spike-and-slab prior (Mitchell & Beauchamp, 1988), the Bayesian Lasso (Park & Casella, 2008), or the horseshoe prior (Carvalho *et al*., 2009) may be cited. In addition, the Bayesian framework also enables the simultaneous integration of multilocus relationships and the data structure such as temporal dependencies. Despite their potential, only few studies explored the application of Bayesian variable selection approaches in the analysis of dynamic phenotypes represented as time series (Heuclin *et al*. 2021, Qu *et al*., 2022). Heuclin *et al*. (2021) introduced the Bayesian Varying Coefficient Model (BVCM), a multilocus model specifically designed for analyzing dynamic phenotypes. It consisted in a multivariate linear model that simultaneously allowed for the selection of genetic markers involved in the variability of phenotypes over time and the estimation of their dynamic effects. These effects were estimated using either interpolation methods, such as B-splines or P-splines, or direct approaches like the random walk process. Marker selection is achieved using a spike-and-slab prior, which yields a posterior probability of selection for each marker, computed simultaneously.

In this study, to demonstrate the reliability and efficiency of the BVCM approach, we applied it to time series phenotypes measured at various time scales (daily, monthly, yearly) across a diverse set of biological species. We included in this study: i) *Saccharomyces cerevisiae* as a yeast model organism with well-characterized dynamic phenotypes, ii) *Fusarium graminearum* as a fungal phytopathogen model for studying host plant interaction dynamics throughout infection cycle, iii) eucalyptus (*Eucalyptus urophylla x E.grandis*) as a fast-growing tree species highly studied in quantitative genetics, and iv) sweet cherry tree (*Prunus avium*) as a fruit tree exhibiting phenological trajectories influenced by diverse environmental pressures. Studying such a panel of organisms has two main objectives: first, to demonstrate the robustness and broad applicability of the BVCM model across a wide range of biological systems; and second, to better understand the genetic architecture of the time related phenotypic plasticity. To do so, the results of the BVCM have been compared with those obtained with a validated method of marker-trait association. The results of the BVCM have been explored deeper by estimating the gain in phenotypic variance explained. Then, the contribution of each genetic region identified to phenotype variability have been evaluated over time.

## 2. Material and Method

### 2.1. Data sets

Independent datasets, produced by distinct research teams participating in a collaborative effort, were obtained from genetic studies carried out on contrasted species (yeast, fungi, eucalyptus, and sweet cherry). For each data set, time series phenotypes were extracted and are briefly described below:

- Yeast data were issued from Peltier *et al*. (2018b) analysing fermentation dynamics of a F1 *Saccharomyces cerevisiae* cross generated by Huang *et al*. (2014). The progeny was composed by 95 genotypes and characterised for 8,070 SNPs homogeneously distributed on all chromosomes (Roncoroni, 2014). In a wine culture media, CO_2_ release was measured by monitoring the fermenters weight loss which represented a time series of 200 time points (Peltier *et al*., 2018a). To limit the complexity of the data set, only one over 20 time points were considered (time series of 10 time points).
- Fungal data were obtained from Laurent *et al*. (2021) and the genetic material described by Laurent *et al.*, (2018). The progeny was composed by 94 F1-strains and genotyped with 483 SNPs markers evenly distributed along the genome. Two time series phenotypes were considered : i) the *in planta* disease severity ratio measured as the number of infected spikelet per spike for 8 time points on two bread wheat cultivars (*Triticum aestivum*), ii) the *in vitro* radial growth assays, measuring the fungal surface at 6 time points thanks to an image analysis automated.
- Eucalyptus data developed by Bartholomé (2014), consist of growth phenotypes and molecular data for 960 full-sibs of an interspecific cross *E. urophylla* × *E. grandis*. A pseudo-test cross method was used to obtain the two parental genetic maps (reference) including 1,429 SNPs for the parent *E. grandis* and 1,323 SNPs for *E. urophylla*. Trunk diameter at 20 cm above ground was measured at 3, 10, 15, 18, 22, 27, 30, and 35 months.
- Sweet cherry data were acquired by Branchereau *et al*. (2023) on a F1 progeny from a cross between two *Prunus avium* cultivars ‘Regina’ × ‘Lapins’. The genetic maps of the parents, produced also with pseudo-test cross method, ‘Regina’ and ‘Lapins’ were composed of 136 SNPs and 124 SNPs respectively. The phenotypes measured were the beginning of flowering date (BF, 10% of the floral buds reached full bloom) and full flowering date (FF, 75% of the floral buds reached full bloom). The phenotyping of 115 genotypes was realized annually between 2018 and 2021 in Forli, Italy.

### 2.2. Heritability estimation

The narrow sense heritability (h^2^) was computed as:

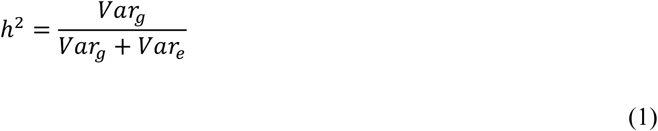

Where *Var_g_* is the additive genetic variance and *Var_g_* + *Var_e_* corresponds to the total phenotypic variance, *Var_e_* being the residual variance. The genetic and phenotypic variances were estimated with the BGLR_1.1.3 package (Pérez & Campos, 2014) using 5,000 iterations and 2,000 burn-in samples and computing the genomic relationship matrix G (VanRaden, 2008) defined by:

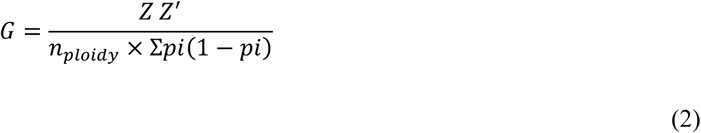

Where *Z* is the centered matrix of genotypes, whose elements are calculated by subtracting the average allele frequency and *n_ploidy_* being the ploidy level of the species. For an individual *j* and a marker *i*, the element *Z_ij_* is defined by:

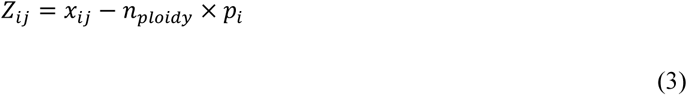

Where *x_ij_* represents the genotype value for individual *j* at marker *i*, and *p_i_* is the allele frequency for marker *i*.

### 2.3. Statistical strategies for quantitative trait analysis and QTL detection

For each data set, the missing data at the phenotypic level were imputed using the missMDA package V1.19 (Josse & Husson, 2016). The best number of components was optimized with the “Kfold” method allowing controlling for overfitting. For molecular data, the missing data were imputed with beagle 5.2 (Browning *et al*., 2018) using default parameters.

The genetic regions involved in the determinism of dynamics phenotypes was explored through two statistical approaches (BLINK and BVCM). Both approaches were based on a multivariate variable selection method, incorporating all markers in the analysis with BVCM and a partial selection of markers with BLINK.

#### 2.3.1 *BLINK* approach (Huang *et al*., 2018)

BLINK method, integrated in GAPIT (Wang & Zhang, 2021), performs a stepwise selection using two multivariate regression models. At each iteration, all markers are tested one at a time in a linear model with a set of pseudo quantitative trait nucleotides (QTN) as covariates initialized as an empty set. The markers are then ranked according to their p-value and filtered for their significance (p-value < 0.01) and only one significant marker within pairwise correlation groups (> 0.7) is retained before evaluation. Then, the best subset of markers is evaluated through Bayesian information criterion. This set is then considered as the new pseudo QTN set. The process is repeated until the set of pseudo QTN remains stable. As a result, the p-values of the set of stable pseudo QTNs are returned and significant QTL are selected after applying the p-values Bonferroni correction with a significance threshold set at 0.05/*n* with *n* being the number of markers used.

BLINK model showed strong performances in the detection of reliable loci in marker-trait association studies. These association studies were carried out for each data set at each time step of the time series. The minor allele frequency was set on 0.05 and three principal components were used to consider the population structure which was negligible in the context of progenies populations. The marker selection threshold was estimated with a Bonferroni correction as:

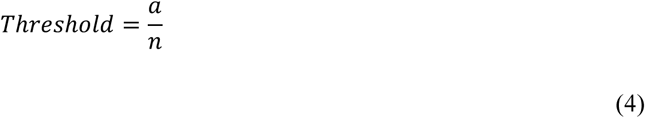

Where *a* is the type I error and *n* is the number of SNPs considered. Manhattan plots and QQ plot were computed with the CMplot_4.5.1 function of the rMVP tool (Yin *et al*., 2021).

#### 2.3.2 *BVCM* approach (Heuclin *et al*., 2021)

The BVCM approach allows analysing all measures of the time series in a single step. In a Bayesian framework, the approach proposes a molecular marker selection method based on a Bayesian spike-and-slab prior combined with a dynamic effect estimation for each marker using different interpolation methods implemented in the model: random walk, Legendre, B-spline or P-spline. Here, the B-spline was used for its simplicity (no hyper parameter settings need to be specified), its capacity to analyze a large number of time steps and because it requires estimating only a small number of parameters.

The model was inferred with a Monte Carlo Markov chains (MCMC). Highly correlated markers were sorted out as described in Heuclin *et al*. (2021) according to a threshold estimated separately for each data set with the aim to limit the complexity of detecting significantly associated genetic regions by summarizing them into representative markers. The correlation threshold and the number of markers retained are indicated in the Table S1.

The identification of relevant markers using the BVCM has been conducted in two steps in order to optimize the results reliability:

- 1^st^ step: the MCMC algorithm was run for 10,000 iterations after a burn-in of 20,000 iterations, a thinning of 5, and for 100 repetitions. The chain convergence was checked by plotting the trace plot of the residual variance (as a proxy of global convergence). The MCMC samples were compiled and the mean of posterior marginal probabilities of variable inclusion were computed. The markers with the highest marginal probabilities were selected to define a subset which was used in the second round. The threshold of selection and the number of markers are indicated in the Table S1.
- 2^nd^ step: the marker subset was used to run again the MCMC algorithm for 10,000 iterations after a burn-in of 20,000 iterations, a thinning of 5, and for 100 repetitions. As in the first step, the chain convergence was checked by plotting the trace plot of the residual variance. The MCMC samples were compiled and the mean of posterior marginal probabilities of variable inclusion were computed. Two thresholds were used to select the significant markers according to their inclusion marginal probabilities. The first threshold was 0.8 if a set of markers was available at this threshold. If not, the second threshold of 0.4 was considered to select significant markers (Table S1).

### 2.4. From cluster to individual marker effect

After QTL detection by each statistical model, the explained phenotypic variance has been explored at the cluster or individual marker levels. Clusters consist of the complete set of significantly associated markers identified by each approach. When evaluating the cluster effect, the corresponding set of markers is assessed.

#### 2.4.1 Cluster effect evaluation

The variance explained by the cluster was estimated with a linear model:

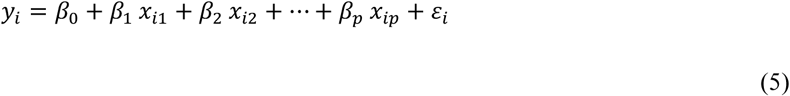

Where y_i_ is the phenotype of the observation *i*, *β_0_* is the model intercept, *β_k_* is the coefficient associated with the *k^th^* variable (or marker), *x_ik_* is the value of the *k^th^* variable for observation *i*, and *ε_i_* is the residual error for the observation *i*. The model was computed with the lme4_1.1-35.5 package (Bates *et al*., 2015) and the R^2^ of the model was used to evaluate the cluster variance explained.

#### 2.4.2 Individual marker effect evaluation

In order to evaluate the variance explained by each marker selected, the same linear model was computed with the relaimpo_2.2-7 package (Groemping, 2007). The R^2^ was decomposed in order to evaluate the phenotypic variance explained by each marker considering collinearity in the model. The marginal contribution of each marker was considered with the Lindeman, Merenda and Gold (LMG) method.

## 3. Results

### 3.1. The high diversity of biological contexts

The data sets which covered a large diversity (Table 1): the progeny size (from 87 to 960 genotypes), the type of trait, the number of measurements (from 4 to 10 steps in time series), and the number of markers available (from 123 SNP to 8,070 SNPs). Phenotypic variabilities were observed both within and between time steps (Fig. S1). The standard deviation varied over the time series and was not homogeneous between species and traits. This statement can be illustrated by the variability of flowering dates of *P. avium* and the growing trait of *E. grandis* and *E. urophylla* that were more stable over time compared to the variability of the CO_2_ release of *S. cerevisiae* and the disease severity and growth of *F. graminearum*.

**Table 1:**
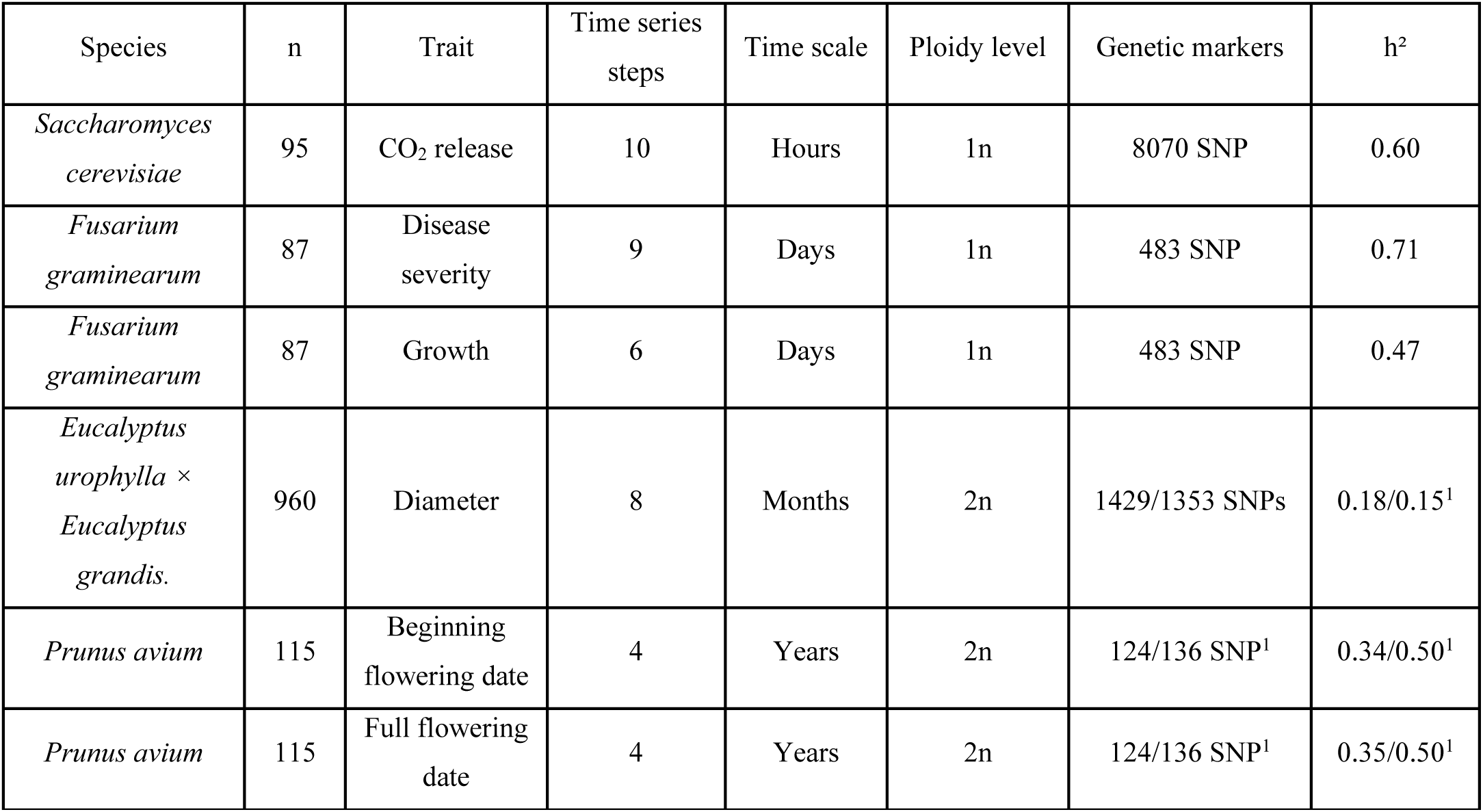
Summary of data sets analysed in the study. The h^2^ column indicates the average narrow sense heritabilities of the different traits over the time series. ^1^ Narrow sense heritability calculated according to each of two the parental genetic matrix.

The h^2^ variability depended on the time series, the species, and the traits considered (Fig. 1). The h^2^ varied from low level for *Eucalyptus* (0.15 to 0.18) to high level for *F. graminearum* disease severity measurement (0.76). The h^2^ varied more largely over time for the CO_2_ release of *S. cerevisiae* compared to flowering dates in *P. avium*, which indicated variability in the genetic contribution of phenotype variability over time. Within the same species, the h^2^ was quite stable over time and trait except for the *F. graminearum* data set in which the h^2^ of disease severity was more stable compared to the h^2^ of growth over time.

**Figure 1:**
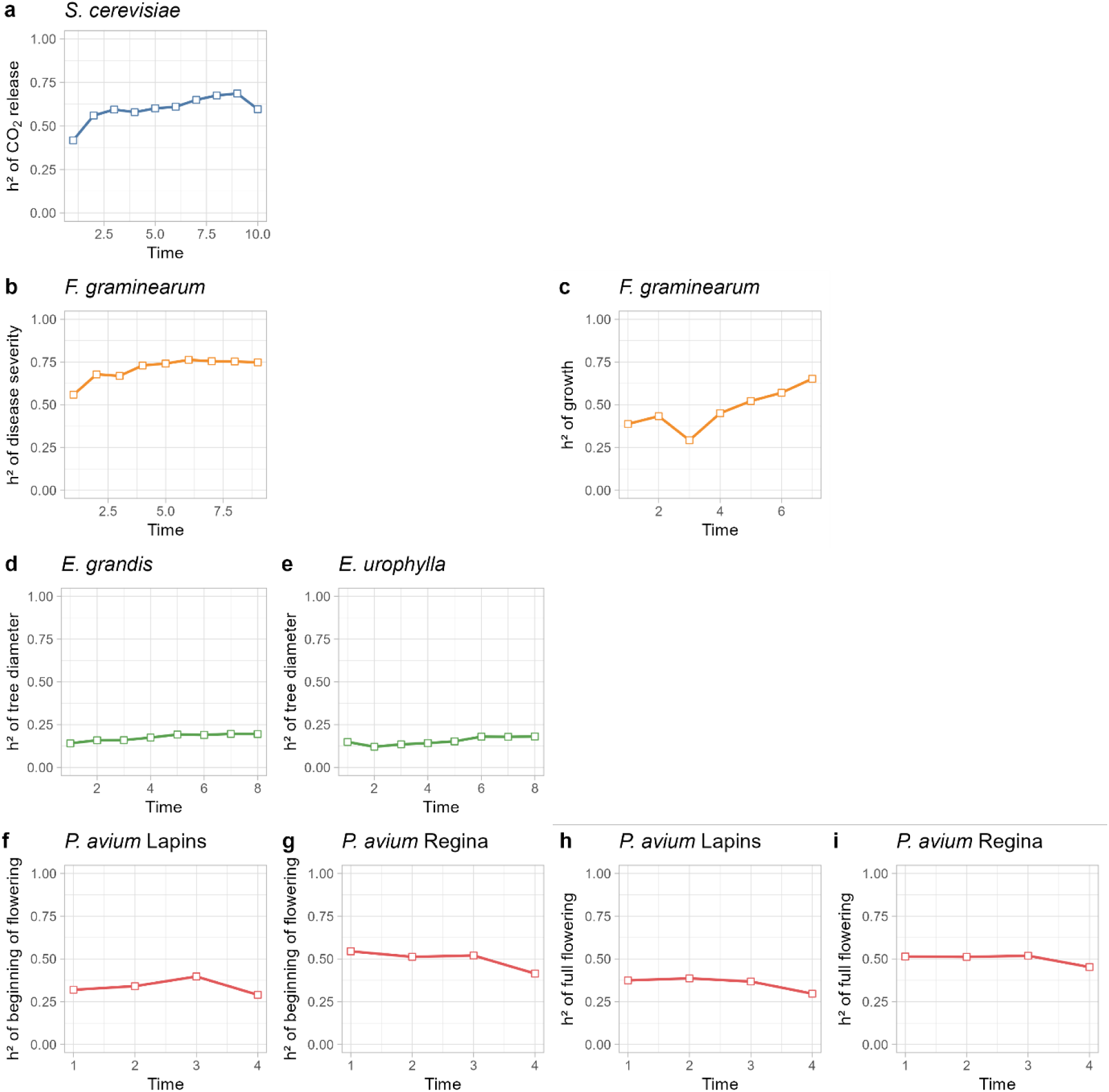
Narrow sense heritability over time. The narrow sense heritability was based on genomic and phenotypic variance and calculated with the BGLR_1.1.3 package (Pérez & Campos, 2014). The genomic variance was obtained from a variance-covariance VanRaden matrix. The data sets consisted of *Saccharomyces* progeny for the CO_2_ release over time (blue, a), in a *Fusarium* progeny (orange) for aggressiveness (b) and growth (c) time series, in an *Eucalyptus* progeny (green) for tree diameter (d and e) time series, and in a *Prunus avium* progeny (red) for the beginning of flowering (f and g) date and the full flowering date (h and i) over years on the base of two parental genetic maps: *Prunus avium* ‘Lapins’ (f and h) and *Prunus avium* ‘Regina’ (g and i).

These different biological contexts offered the opportunity to test a same methodological approach to identify underlying genetic determinants in a panel of time related phenotypic plasticity.

### 3.2. Increasing the QTL detection power

In this section, the results of the two QTL selection approaches (BLINK and BVCM) are presented. The phenotypic variance explained (PVE) indicates the part of the phenotypic variance that can be attributed to an individual genetic marker or a set of markers. The PVE reported corresponds to the average proportion of PVE contributed by each individual marker over the considered time series. Within each approach, some of the detected markers overlapped. However, since their PVE was estimated within distinct marker clusters, the resulting PVE values could differ for the same marker between the two approaches.

#### 3.2.1. BLINK approach detected major genetic variants

The BLINK analysis identified three genetic regions significantly associated with CO_2_ release across the ten-time steps of the *S. cerevisiae* time series (Fig. 2). The associated markers were located on three different chromosomes: Chr4 (Sc4_852909), Chr5 (Sc5_305580), and Chr16 (Sc16_878259). Additionally, a fourth genetic region on Chr5 (Sc5_462977) was close to the significance threshold and considered associated with the variability in CO_2_ release. Over the time series, these markers accounted for a PVE ranging from 7.2% (Sc5_462977) to 16.8% (Sc16_878259). Over the course of the time series, marker Sc4_852909 was identified at three time points, Sc5_305580 at four time points, and Sc16_878259 at seven time points.

**Figure 2:**
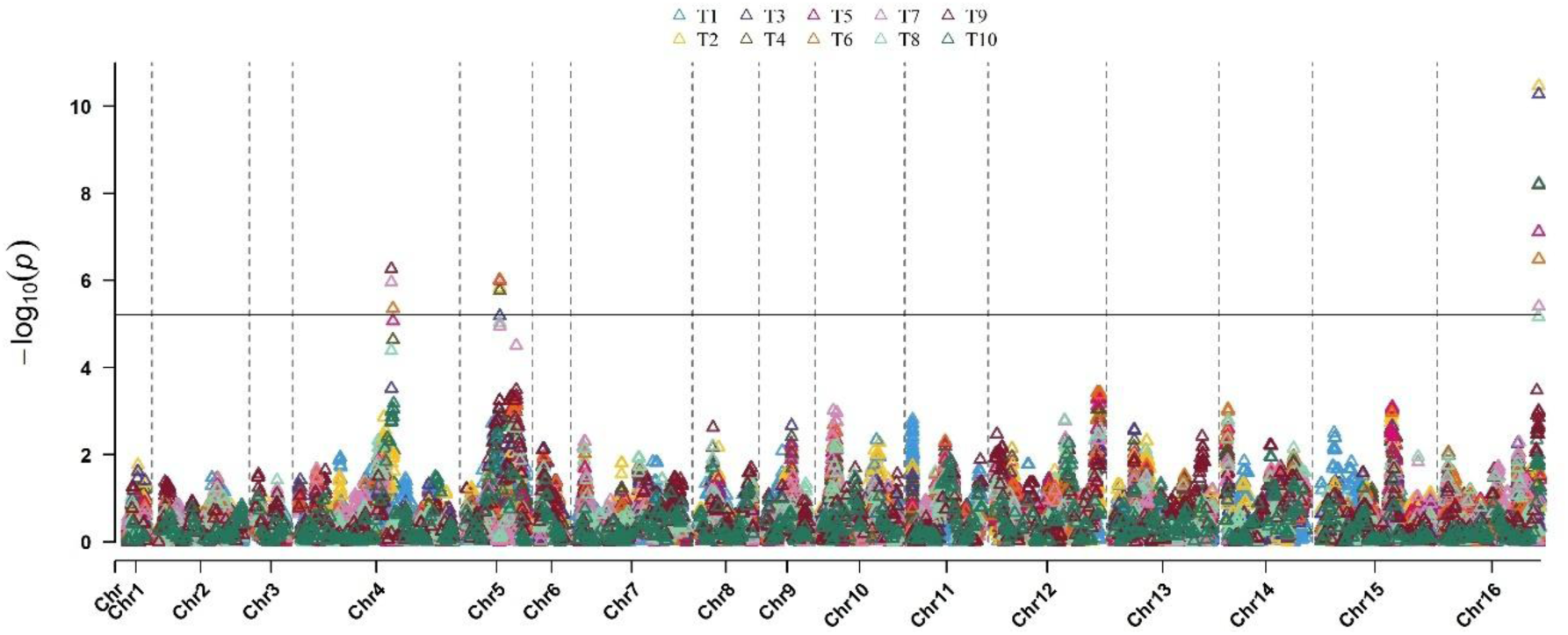
Multi-traits Manhattan plot of marker-trait association results for the 10 time points of the CO_2_ release time series of the *S. cerevisiae* data set. The results are issued from the BLINK model and each time step of the time series is represented by a specific color.

Two genetic regions were significantly associated with the disease severity across the nine time points of the *F. graminearum* time series (Fig. S2). The associated markers were located on Chr1 (Fg1_92) and Chr2 (Fg2_115). Over the time series, these markers accounted for 81.3% and 1.9% of the PVE respectively. Marker Fg1_92 was identified for all time steps of the time series whereas marker Fg2_115 was only identified for the first time step. The genetic region surrounding Fg1_92 was also associated with fungal growth across the seven time points of the time series (Fig. S3). The two neighboring markers Fg1_92 and Fg1_93 were each identified for three time steps of the fungal growth. These two markers represented the same genetic region and were never identified together in the BLINK analysis. Over the time series, the genetic region accounted for 36.9% of the PVE.

The genetic regions significantly associated with the diameter across the eight-time steps of the *Eucalyptus* time series were analyzed for each parental genetic map separately (*E. grandis* and *E. urophylla*). The BLINK analysis identified two genetic regions from the *E. grandis* genetic map located on two chromosomes: Chr2 (Eg2_42.04) and Chr5 (Eg5_50.45) (Fig. S4). These markers accounted for 1.5% and 1.0% of the PVE respectively. Over the course of the time series, marker Eg2_42.04 was identified at one time step and marker Eg5_50.45 was identified at three-time steps. One genetic region was identified from the *E. urophylla* genetic map located on Chr4 which was supported by four genetic markers (Eu4_50.76, Eu4_51.35, Eu4_53.45, Eu4_53.54) for one time point over the time series (Fig. S5). These markers accounted for a similar PVE of 0.4%. Eu4_51.35 accounted for a slightly higher PVE and was selected to represent the genetic region which then accounted 1.5% of the PVE.

The genetic regions significantly associated with BF and FF across the four time points of the *Prunus avium* time series were analyzed for each parental genetic map separately (*P. avium* ‘Lapins’ and *P. avium* ‘Regina’). For the BF time series, the BLINK analysis identified one genetic region from the *P.avium* ‘Lapins’ genetic map located on Chr5 (Pl5_62423) (Fig. S6). Pl5_62423 accounted for 5.0% of the PVE and was identified for one time step. One genetic region was identified from the *P. avium* ‘Regina’ genetic map for one time step of the BF time series (Fig. S7). The marker identified was located on Chr4 (Pr4_29266) and accounted for 21.5% of the PVE. No marker was identified from the *P. avium* ‘Lapins’ genetic map for the FF time series (Fig. S8) while the marker Pr4_29266 was also identified from the *P. avium* ‘Regina’ genetic map for one time step the FF time series (Fig. S9) and explained 19.4% of the PVE.

#### 3.2.2. BVCM approach detected additional weak effect genetic variants

The BVCM identified 16 genetic markers associated with CO₂ release across the ten time steps of the *S. cerevisiae* time series (Fig. 3). Among these, four markers (Sc4_881976, Sc5_305580, Sc5_462977, and Sc16_878259) were located in close proximity to those detected by BLINK, and were thus considered to be common to both approaches. These markers individually accounted from 5.6% (Sc4_881976) to 12.9% (Sc16_878259) of the PVE. Additionally, 12 markers (Sc1_95310, Sc7_65336, Sc8_130148, Sc8_180363, Sc8_476586, Sc10_112742, Sc11_363180, Sc11_639434, Sc12_240970, Sc12_659981, Sc14_27762, and Sc15_712830) were uniquely identified by the BVCM and accounted from 0.6% (Sc12_240970) to 5.6% (Sc11_363180) of the PVE.

**Figure 3:**
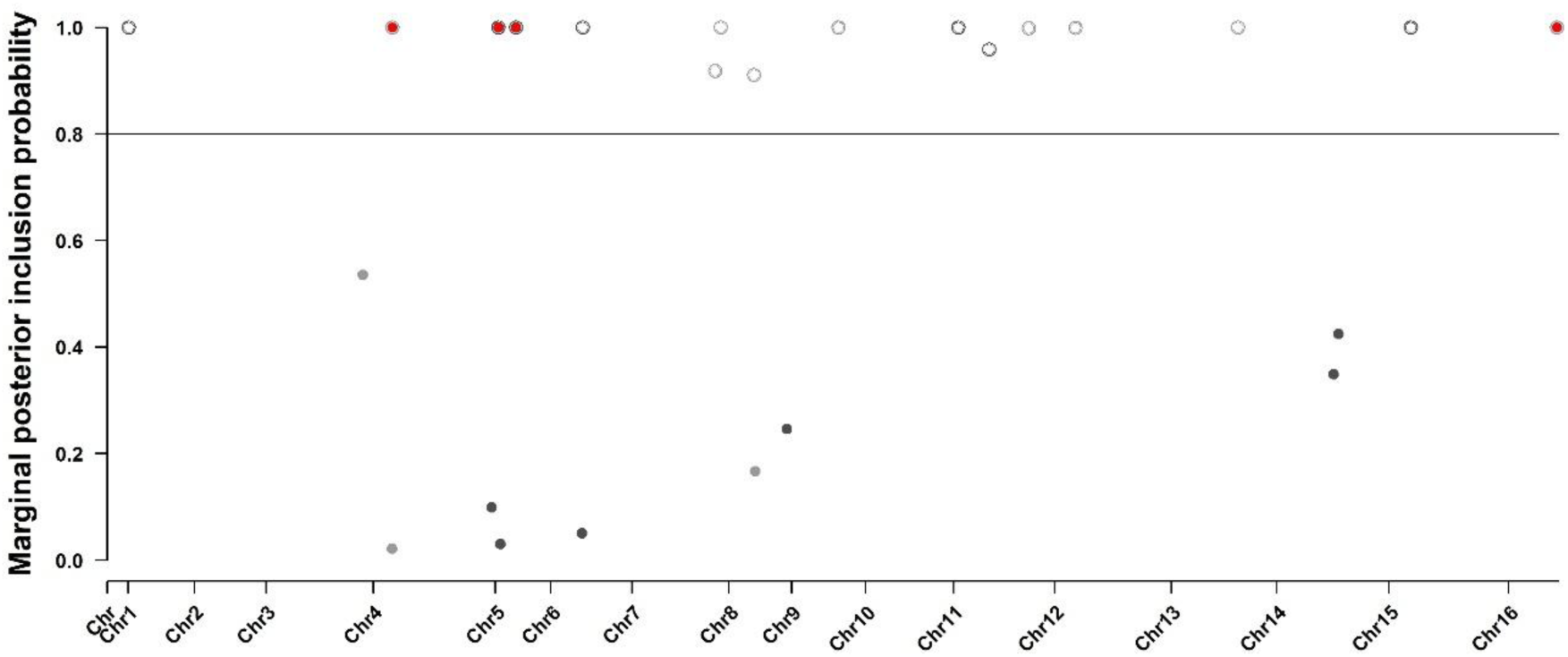
Marginal posterior inclusion probabilities of the *S. cerevisiae* genetic markers using the Bayesian Varying Coefficient Model. The probability was calculated according to the association between each marker and the variability of the CO_2_ release time series. The line represents the selected threshold for marker selection and circles indicate the selected marker. Markers corresponding to genetic region also detected by the association mapping approach are filled in red.

One genetic marker was associated with the disease severity across the nine-time steps of the *F. graminearum* time series (Fig. S10). This marker was located on Chr1 (Fg1_91) and accounted for 77.3% of the PVE. The marker belonged to a genetic region detected through the BLINK approach with the marker Fg1_92. The BVCM further identified nine genetic markers associated with the growth time series (Fig. S11). They include the genetic region identified by BLINK on Chr1 (Fg1_91) which accounted for 18.7% of the PVE. The additional markers identified with BVCM were located on Chr1 (Fg1_86, Fg1_187, Fg1_212, Fg1_231), Chr2 (Fg2_398), Chr3 (Fg3_24, Fg3_172), and Chr4 (Fg4_219) and accounted from 0.6% (Fg3_24) to 16.4% (Fg1_86) of the PVE.

Six genetic markers were associated with the tree diameter across the eight-time steps of the *Eucalyptus* time series from the *E. grandis* parental genetic map (Fig. S12). Among them, two markers located on Chr2 (Eg2_42.04) and Chr5 (Eg5_50.45) were also identified with BLINK and accounted for 1.5% and 0.9% of the PVE respectively. Four markers were additionally identified with BVCM on Chr3 (Eg3_78.55), Chr8 (Eg8_15.87, Eg8_69.17), and Chr10 (Eg10_19.14) and accounted from 0.5% (Eg8_69.17) to 1.3% (Eg3_78.55) of the PVE. Three markers were identified from the *E. urophylla* parental genetic map (Fig. S13). One marker was also identified with BLINK on Chr4 (Eu4_51.35) which accounted for 1.3% of the PVE. Two markers were additionally identified with BVCM on Chr3 (Eu3_54.27) and Chr4 (Eu4_7.1) and both accounted for 0.8% of the PVE.

Based on the BF time series measured on *Prunus*, three genetic markers from the Lapins parental map were associated with the time series (Fig. S14). One marker located on Chr5 (Pl5_62423) was also identified with BLINK accounting for 3.3% of the PVE. Two markers located on Chr1 (Pl1_119926) and Chr5 (Pl5_57602) were additionally identified with the BVCM accounting for 8.0% and 2.2% of the PVE respectively. Based on the ‘Regina’ genetic parental map, two genetic marker were associated with BF time series (Fig. S15). One marker was located on Chr4 (Pr4_29266), accounted for 20.6% of the PVE, and was also identified with BLINK. The second was additionally identified with the BVCM and located on Chr5 (Pr5_14458) and accounted for 5.0% of the PVE. For the FF time series of *Prunus*, two markers from the Lapins parental map located on Chr1 (Pl1_161670) and Chr6 (Pl6_8214) were associated with the trait over time (Fig. S16). These markers were only identified with BVCM and accounted for 9.9% and 4.7% of the PVE respectively. From the ‘Regina’ genetic parental map, one marker was associated with the FF time series (Fig. S17). The marker was located on Chr4 (Pr4_29266) and was also identified with BLINK. This marker accounted for 19.4% of the PVE.

### 3.3. Clusters of markers identified by BVCM increases the phenotypic variance explained

The PVE was evaluated for the cluster of markers identified with the BLINK and the BVCM approach separately (Fig. 4). The clusters identified with BVCM increased the PVE compared to those identified with BLINK except in two contexts described below. On average, the clusters identified with BVCM increased the PVE by 7% compared to clusters identified by BLINK. The higher increase of PVE was observed for the CO_2_ release by *S. cerevisiae* for which the BVCM cluster was on average 27.3% higher than the BLINK cluster across the time series.

**Figure 4:**
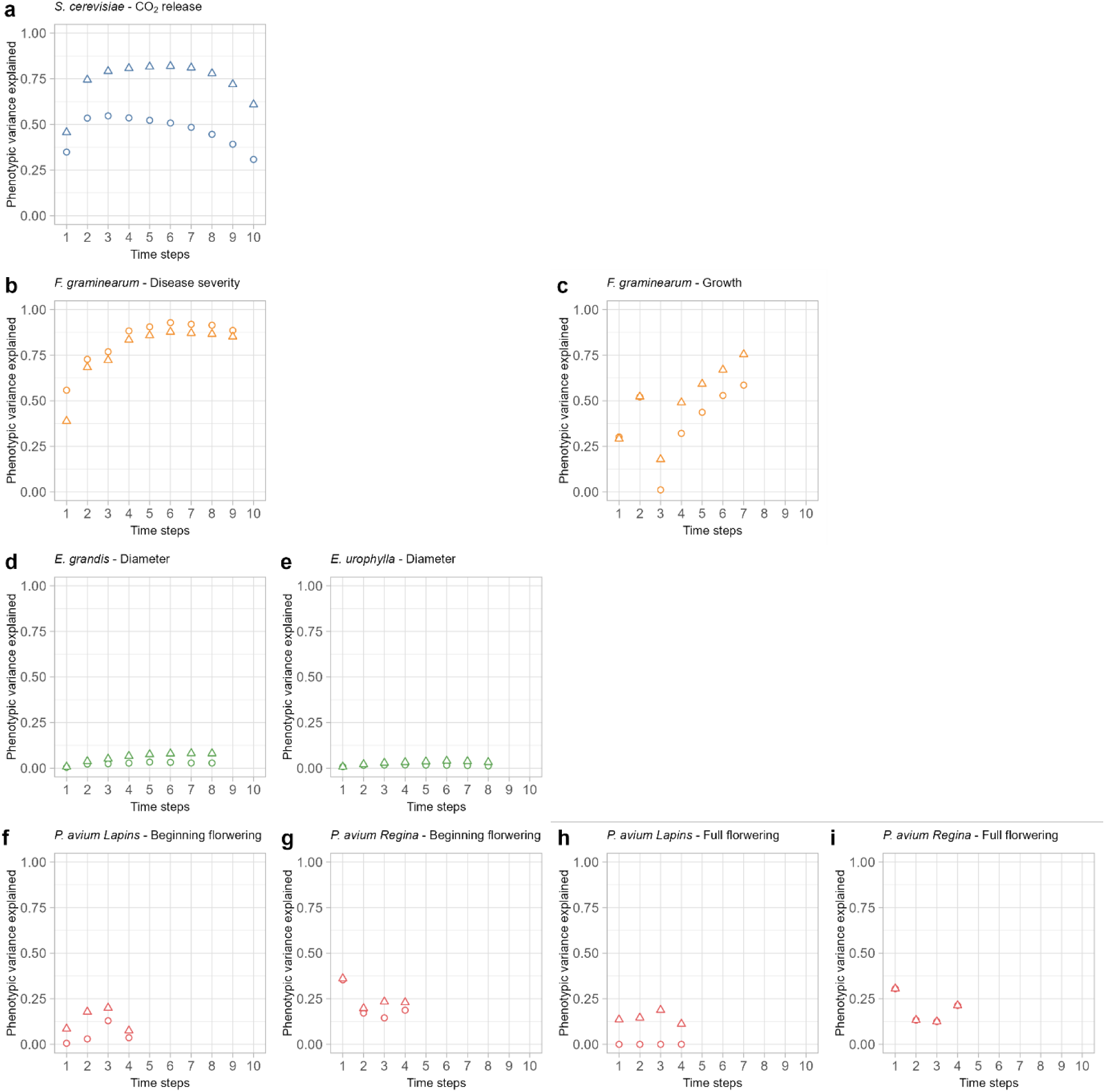
Phenotypic variance explained by the marker sets identified by BLINK (circle points) and BVCM (triangle points) over time. The data sets have been obtained in a *Saccharomyces* progeny (blue) for the CO_2_ release over time (blue, a), in a *Fusarium* progeny (orange) for aggressiveness (b) and growth (c) time series, in an *Eucalyptus* progeny (green) for diameter (d) and height (e) time series, and in a *Prunus avium* progeny (red) for the beginning of flowering date and the full flowering date over years on the base of two parental genetic maps: *Prunus avium* ‘Lapins’ (f and g respectively) and *Prunus avium* ‘Regina’ (h and i respectively).

The clusters identified with BVCM showed lower or equal PVE in two contexts. Firstly, the disease severity time series of *F. graminearum* was associated with the marker Fg1_92 with BLINK and Fg1_91 with BVCM. The marker Fg1_92 allowed to obtain a higher PVE for the all-time steps compared to FG1_91 (4% higher on average). Moreover, the marker Fg2_115 only highlighted by BLINK for the first-time step of the time series led to an increase of the PVE for the BLINK cluster compared to the BVCM cluster for the first-time step (55.8% and 38.9% respectively). Secondly, the full flowering date time series of *P. avium* allowed identifying the same genetic marker (Pr4_29266) with the two approaches from the ‘Regina’ parental map, leading to equal PVE across the time series.

### 3.4. Markers individual effects revealed weak or transitory genetic effects

In all the studied organisms, the markers commonly identified by the two approaches were those with the highest PVE for each time series. The additional markers identified by the BVCM explained a lower phenotypic variance or were more time-specific.

As illustrated for the *S. cerevisiae* (Fig 5), markers identified by both BLINK and BVCM showed the highest PVE over the CO_2_ release time series. The markers commonly selected by the two methods have shown a higher maximum for the PVE over time (from 8.2% for Sc4_881976 to 21.9% for Sc16_878259) compared to the BVCM additional markers whose PVE patterns evolved between 0% and 7% over the time series. The PVE patterns of common makers where similar when evaluated in the two clusters, however, the PVE was lower in the BVCM context for each time steps of the time series. While common markers showed similar pattern to the h^2^ of the time series, additional markers have shown specific patterns increasing at the end of the time series such as Sc8_130148 and Sc8_476586.

**Figure 5:**
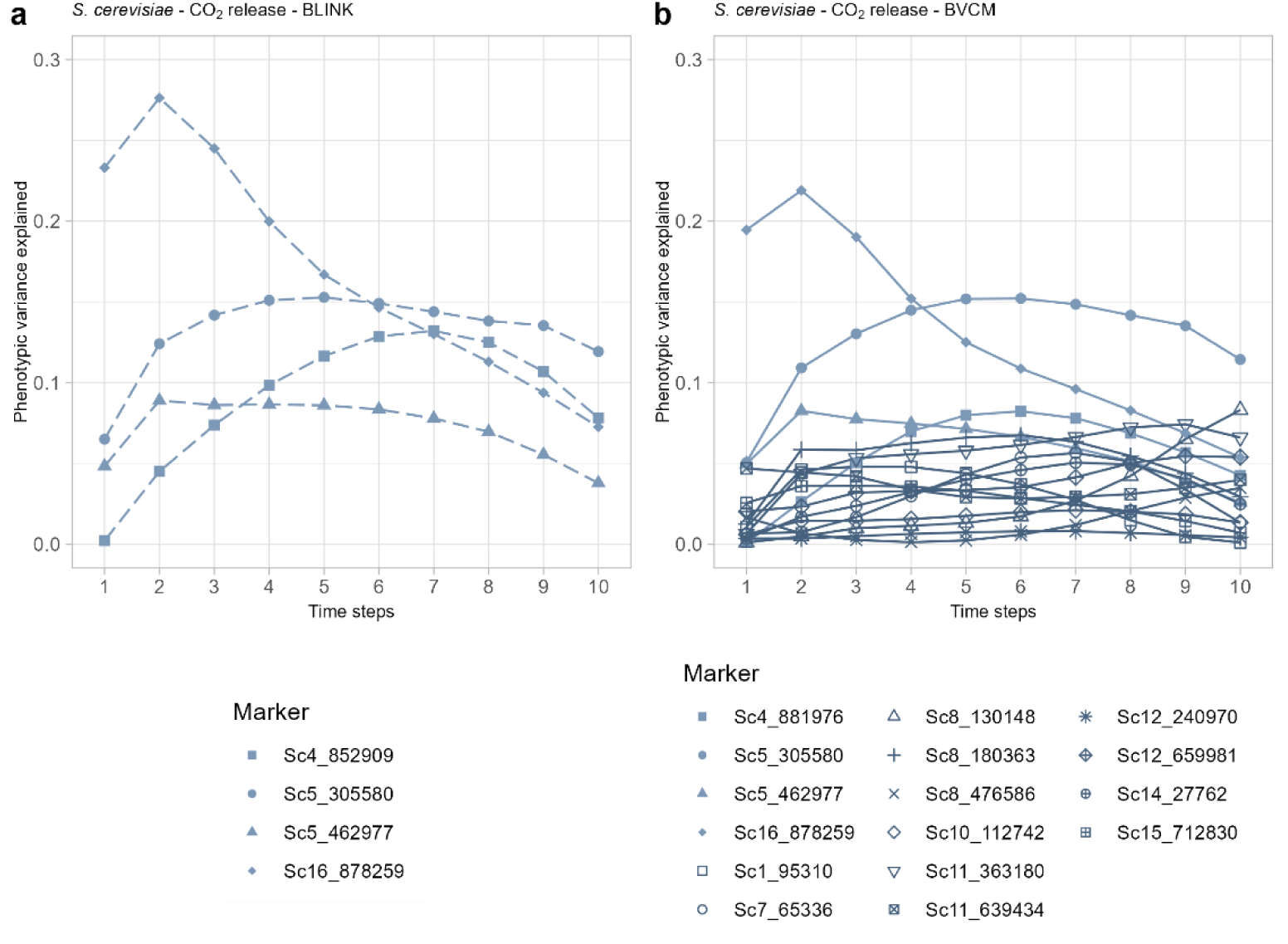
Phenotypic variance explained over time for the CO_2_ release time series of *S. cerevisiae* by markers selected by the BLINK (dotted lines, a) and the BVCM (full lines, b) approaches. The markers commonly identified by BLINK and BVCM are indicated by light colored lines in B. Each marker is represented by a different point shape.

Considering the *F. graminearum* data set, the markers identified for the disease severity on Chr1 by BLINK and BVCM (Fg1_92 and Fg1_91 respectively) explained the same part of the PVE over time (Fig. S18). The additional marker identified by BLINK (Fg2_115) explained a weak part of the PVE for the first step of the time series and was null for the other time steps. The markers identified by BLINK and BVCM for the growth dynamics of *F. graminearum* were similar in term of position and PVE over the time series (Fig. S19). The BVCM selected additional marker which explained a lower part of the PVE over time. These markers explained an increasing part of the PVE as the time series advanced.

The markers identified by BLINK and BVCM for the diameter dynamics of *E. grandis* explained a small part of the PVE over time (from 0.1% to 2.2%) (Fig. S20). The BVCM specifically identified four additional genetic markers which explained lower PVE for the first steps of the time series compared to one of the two common markers but they explained a higher PVE as the time series progressed. The marker commonly identified by BLINK and BVCM for the diameter dynamics of *E. urophylla* explained from 0.7% to 1.5% of PVE over time (Fig. S21). The additional marker identified by BLINK explained lower part of the PVE from 0% to 1.4% over time.

Based on the beginning of flowering time series of the *P. avium* data set, Pl5_62423 was identified by BLINK and BVCM from the ‘Lapins’ genetic parental map for (Fig. S22). BVCM additionally identified Pl1_119926 and Pl5_57602. Pl5_57602 showed a PVE pattern similar to Pl5_62423. Pl1_119926 was more associated with the second time step of the time series. The marker selected from the *P. avium* ‘Regina’ parental genetic map by BLINK and BVCM for the beginning of flowering date (Fig. S23) Pr4_29266. The additional marker Pr5_14458 identified by the BVCM has shown opposite pattern of PVE compared with Pr4_29266.

For the full flowering time series, while no marker was identified with BLINK, two markers were identified by BVCM from the ‘Lapins’ genetic parental map with different PVE patterns (Fig. S24). The marker Pl1_161670 caught from 4.8% to 13.8% of the PVE and had the maximum effect at the second time point of the time series. The marker Pl6_8214 explained from 0.8% to 8.8% of the PVE with a maximum effect for the first time point. The marker selected from the *P. avium* ‘Regina’ parental genetic map by BLINK and BVCM for the full flowering date was Pr4_29266 (Fig. S25). This marker has shown similar PVE patterns for the two time series.

## 4. Discussion

Contrasted datasets were used to assess the performances and the advantages of the recently proposed BVCM model for analyzing the genetic determinism of time-series phenotypes for multiple traits/species. The BLINK method, known for reducing false discovery rates in GWAS (Huang *et al*., 2018), was applied in a time by time approach for each time series. BLINK results were then compared with those obtained with BVCM. Across these different contexts, BVCM successfully identified major QTLs already detected by BLINK while also uncovering additional QTLs with more punctual or moderate effects over time. These newly identified QTLs allowed to increase the genetic PVE. The robustness and broad applicability of BVCM make it particularly well-suited for detecting QTLs associated with dynamic phenotypes, taking advantage of a multilocus approach.

### 4.1. Highly multidimensional analysis to increase the QTL detection power

The BVCM approach identified larger sets of markers compared to the BLINK stepwise approach. For almost all species, the markers selected by the stepwise approach were included in these sets and corresponded to markers with the highest effects. These results confirmed the ability of BCVM to identify additional markers with potentially biological relevance that can be attributed to two key factors.

First, the analysis of all time points, while accounting for the underlying temporal structure, enhanced the statistical power by increasing the number of observations relative to the number of parameters. This is especially useful when narrow-sense heritability evolves over time (Qu *et al*., 2022). Indeed, as observed in *S. cerevisiae* and in *F. graminearum*, the analysis of traits with increasing heritability over time helps to identify/estimate non-null markers with weak effects at the beginning of the follow-up. With the BVCM method, as we are evaluating a time-series phenotype, we can rely on the trajectory of the phenotype and on periods of high h^2^ to detect weak signals that would not have been distinguishable from noise during periods of low h^2^ (Manolio *et al*., 2009). However, we observed that the differences between the two approaches are less pronounced when the temporal structure or/and the narrow-sense heritability are weak as observed for the diameter times series form the Eucalyptus data set.

Second, the simultaneous analysis of all markers using a Bayesian variable selection approach enables the identification of markers with weaker effects (Hoggart *et al*., 2008; Xu *et al*., 2018c; Adhikari *et al*., 2023; Osorio-Guarin *et al*., 2024). Based on the “omnigenic model” of complex traits, despite weak effect variants spread across the genome have no clear function for traits variability, they can impact the regulatory networks of the trait and explain a significant part of h^2^ (Boyle *et al*., 2017). While they can explain a large part of the phenotypic variance, the detection of QTL with weak effects represent a challenge in linkage analysis and association studies because it is hard to statistically distinguish true weak signals from noise (Yang *et al*., 2010; Shi *et al*., 2016; Boyle *et al*., 2017). Thus, a special attention has been paid to weak effect variants in GWAS mainly for human diseases because a high number of them can significantly increase risk factors (Vsevolozhskaya *et al*., 2019). For instance, Wu *et al*. (2014) proposed the HC-methods which allowed them to detect signals for very rare and weak effect variants by considering them in clusters (SNP-set) of linked variants. They were able to distinguish genetic true weak signals from noise associated with Crohn’s disease in GWAS. Vsevolozhskaya *et al*. (2019) proposed a method of Augmented Rank Truncation (ART) in order to analyze the top ranking SNP and select the best set of markers. They were able to select variants that would not have passed the significance threshold with p-value evaluation after GWAS. Other screening methods have also been proposed in order to detect weak effect variants when GWAS approaches lacked power such as false negative control (Jeng *et al*., 2024). Here, because we calculated the posterior marginal probabilities of association in the Bayesian context, we could select the most appropriate threshold for variants consideration and detect weak effect variants without supplemental statistical evaluation.

Our study also demonstrated that the Bayesian approaches, commonly used in genomic prediction, represent powerful alternatives in QTL mapping with a great flexibility in terms of integrating structure from various sources, such as known genetic relationships or temporal phenotype structures through variance covariance matrix (Heuclin *et al*., 2021; Qu *et al*., 2022).

### 4.2. Two-steps to improve statistical accuracy

In this study, we proposed a two-step approach for identifying relevant markers. The first step, which consisted in running 100 MCMC algorithms, allowed to handle the multi-collinearity within the marker dataset and identified genetic regions with high probability of association. The second step further refined the results by reducing the number of variables. This two-step approach made it possible to reduce the trade-off between the number of parameters and the number of observations, and thus to identify a set of promising genetic markers. Two steps have been used in previous studies using Bayesian models for GWAS. For instance the BGWAS (Bayesian GWAS) model is based on a first step of variables screening in order to select a set of promising genetic markers (Williams *et al*., 2023). A second step selects the more relevant markers in the set. This approach allowed to reduce the false positives detection in BGWAS. Yazdani & Dunson (2015) used a similar procedure with a first screening of promising SNPs by a Bayesian method, each SNP being evaluated individually. The second step consisted in a hierarchical model to simultaneously analyze the effect of all the promising SNP in the set. This method led to higher performances in genomic prediction and reduced the computational cost in a simulated data set. In a different way, Li *et al*. (2011) started by transforming the phenotypic values through supervised PCA in order to catch the overall genetic structure and reduce noise in data. The second step consisted in a hierarchical Bayesian model with a Lasso regression selection method (Tibshirani, 1996). The two steps approach led to an increase in results stability and brought new insights into the genetic architecture of the human body mass index. In the BVCM approach, the use of two steps made it easier to identify genetic markers truly associated with the time series variability. To avoid the need for a preliminary filtering step that removes highly correlated molecular markers, future research could improve the BVCM approach by incorporating both genomic structuration of markers and temporal structuration directly into the selection prior.

### 4.3. Genetic dynamics of phenotype establishment

Compared to the BLINK model, the BVCM combined a multivariate varying coefficient model with Bayesian variable selection approach, allowing to catch fine temporal genetic dynamics. Across the studied contexts, heritabilities exhibited temporal variation over time, with a slight increase in *Eucalyptus* (from 0.14 to 0.20) which was more pronounced in *S. cerevisiae* (from 0.49 to 0.79). The contribution of each genetic region in the determinism of trait dynamics was also variable over time, such as Sc16_878259 of *S. cerevisiae* being able to explain from 5.4% to 21.9% of the CO_2_ release PVE. These observations can be related to different biological processes for which several underlying genetic determinants were involved and occurred in sequence over time. For instance in *Arabidopsis thaliana*, the development and maintenance of certain phenotypes are regulated by the CLAVATA-WUSCHEL pathway, which can either maintain the apical meristem for plant growth or transform the apical meristem into inflorescence meristem for seed production over time and environments (Mayer *et al*., 1998; Schoof *et al*., 2000). In barley, genes have been detected for being associated with specific stages of grain filling (Du *et al*., 2019). In sticklebacks, the ontogeny of the body size is mediated by different genetic regions over stages (Fraimout *et al*., 2022). Thus, some ontogenic stages may represent key shifts in gene expression and cannot be observed in a one-time analyze. Therefore, identifying additional genetic regions that influence phenotype development over time could provide new opportunities for selection in breeding programs. The study of longitudinal data also allowed us to observe h^2^ dynamics over time, which could reveal critical windows of high h^2^ that should be focused to enhance breeding programs.

The BVCM, by integrating the time and genetic structures of data sets, allowed to increase the number of loci detected compared to BLINK. According to the Fisher infinitesimal model (Fisher, 1919), complex quantitative traits are supposed to be controlled by many genetic regions with small effects. By identifying a higher number of genetic regions, we were able to explain a larger part of the h^2^ over time. Because a higher number of loci were considered for estimating the PVE, the PVE calculated for each individual marker decreased when calculated in the BVCM cluster compared to BLINK cluster. It suggests that considering only the markers with the highest genetic effects not only overlooks a part of the genetic variance (Manolio *et al*., 2009) but also leads to overestimating the PVE of selected markers. In addition to the additive variance significantly explained by markers, the genetic architecture of complex traits is expected to account for variants interactions such as epistatic or dominance effects (Holland, 2007; Mackay, 2009). Multilocus approaches can detect clusters of markers with potential epistasis interactions (Bocianowski, 2014) which can have a significant impact on the genetic determinism of complex quantitative traits (Phillips, 2008; Mackay, 2009). By considering all markers in a multivariate model, the multilocus approach would allow to catch such interactions between genetic regions. Moreover, epistatic interactions must be considered for their potential impact on the adaptive value of organisms (Scarcelli *et al*., 2007), for their role in modulating the heterosis effect in hybrids (Sang *et al*., 2022), and for their ability to explain a part of the missing heritability (Slim *et al*., 2020; Balvert *et al*., 2024) by accounting for a non-additive part of the genetic variance (Mäki-Tanila & Hill, 2014). Our approach account for genetic interactions but identifying the genetic regions interacting or evaluating the magnitude of such effects remains challenging to measure and clearly identify (Tyler *et al*., 2009; Hill, 2010; Sakai *et al*., 2021; Ponte-Fernandez *et al*., 2022).

The ability of the BVCM to identify loci with weak effects depended of the data set parameters. The population size, the number of time steps in the time series, the number of genetic markers and the h^2^ of the trait had an effect on the ability to improve the results with the BVCM compared to BLINK. Previously, a comparison between the GBLUP and BayesB revealed difficulties in selecting the best model because each could be superior to the other according to the population size, the number of QTL and the broad sense heritability (h^2^) used in simulated data (Daetwyler *et al*., 2010). Moreover, the accuracy and confidence of QTL detection methods are reduced in the context of low h^2^, small population size, low effect QTL or low density marker (Manolio *et al*., 2009; Wang *et al*., 2012; Mollandin *et al*., 2021). Here, the *Prunus avium* data set had a low number of time steps and markers, which led to a similar number of genetic regions identified with the BVCM and BLINK. In contrast, the *Eucalyptus* data set had a higher number of time steps and individuals, but the BVCM was also unable to identify many new genetic regions associated with the time series. Hence, the high statistical power of this data set and the low genetic variance of the time series (low h^2^) allowed the BLINK approach to catch most of genetic effects. However, even in these cases, the BVCM allowed to assess genetic effect of dynamics QTL over time with transitory, delayed, increasing or decreasing effects, which could be masked in an approach done time by time. These findings contribute to a deeper understanding of the genetic architecture of complex traits, by reducing the issue of missing heritability with a potential in enhancing breeding efficiency (Dwivedi *et al*., 2024).

### 4.4. Time shaping of phenotypic plasticity across diverse organisms

Weak effect variants can represent significant elements of the time related phenotypic plasticity by their contribution to the variance and their interactions (pleiotropy, epistasis, and development) at the genetic level. For instance, Peiffer *et al*. (2014) compared several models for the genetic architecture of maize plant height and the more polygenic (GBLUP) allowed the best prediction for the trait. Still in maize, the flowering time was genetically controlled by a large number of weak effect loci instead of few major QTLs (Buckler *et al*., 2009). Small effect variants were also involved in the genetic architecture of human traits such as human face (Liu *et al*., 2012; Hallgrimsson *et al*., 2014; Adhikari *et al*., 2016). The consideration of traits dynamics in combination with multilocus analysis is meaningful in this context because it increases the opportunity to detect weak effect genetic variants.

Developmental timing is essential for understanding phenotype plasticity. Conith *et al*. (2021) showed in the context of skull bone development in a fish progeny of cichlids that the first bones to form impact the formation of subsequent bones and confuse the relationships between phenotype and genotype. This work also showed that the developmental sequence can be constrained by functional timing because the time related phenotypic plasticity of early forming traits influences the establishment of subsequent traits. The study of branchial bone length variation over time in sticklebacks revealed loci associated with multiple developmental stages whereas the effect of other loci were restricted to juvenile or adult stage (Erickson *et al.*, 2014). It allowed them to better understand the genetic architecture of phenotype establishment and the underlying evolutionary processes. The analysis of the genetic architecture of time series phenotypes, which can be related to functional or developmental plasticity, allowed to understand trait relationships, potential trade-off over time and the underlying genetic dynamics.

The BVCM method experienced on contrasted data sets allowed us to acquire more exhaustive and informative results about the genetic architecture of the time related phenotypic plasticity. We were able to handle the complexity of time and genetic structures in a single statistical approach. The rapid increasing of sequencing and phenotyping technologies lead to the acquisition of larger data sets, with multiple measurements on the same genetic unit. While the high-throughput phenotyping remains a challenge in perennials because of their long-life cycle, the increasing of the number of measurements over years lead also to large data sets which require powerful modeling approaches such as BVCM. By capturing the genetic factors influencing the phenotypic variability across time, our approach will contribute to a deeper understanding of the dynamic nature of genotype–phenotype relationships. More broadly, this work raises the foundation for extended applications in functional genomics, evolutionary ecology, and crop breeding programs, in which temporal plasticity remains crucial for predicting and selecting key quantitative complex traits.

## Acknowledgements

We acknowledge the GPR Bordeaux Plant Sciences for the support provided within the framework of the IdEX Bordeaux University “Investments for the Future” program.

## Statements and Declarations

### Fundings

This study received financial support from the French government in the framework of the IdEX Bordeaux University “Investments for the Future” program / GPR Bordeaux Plant Sciences

### Competing Interests

The authors declare the following financial interests/personal relationships which may be considered as potential competing interests: Philippe Marullo reports financial support was provided by BIOLAFFORT. Other authors declare that they have no known competing financial interests or personal relationships that could have appeared to influence the work reported in this paper.

### Authors Contributions

EM and JM conceptualized the study and obtained fundings. All authors contributed to the set up of the experimental design. Data analysis was performed by LB with the support of BH and MD. The first draft of the manuscript was written by LB, revised and edited by EM and JM and all authors contributed to correct and improve the paper quality. All authors read and approved the final manuscript

### Data Availability

The datasets generated during and/or analysed during the current study are available from the corresponding author on reasonable request.

### Ethics approval

Not applicable

### Consent to participate

Not applicable

### Consent to publish

Not applicable

