## Supplemental files for "A Bayesian multidimensional approach to decipher the genetic basis of dynamic phenotypes in multiple species"

**Supplemental information**

Table S1: Filters applied on data sets in the Bayesian varying coefficient model procedure

| | | | | Pre-processing | | 1st round of BVCM | | 2nd round of BVCM | |
|---|---|---|---|---|---|---|---|---|---|
| Biological species | Initial markers (SNP) | n | Trait | cor threshold | Filtered markers | Threshold 1st round | marker 1st round | Threshold 2nd round | marker 2nb round |
| *Saccharomyces cerevisiae* | 8070 | 95 | CO2 release | 0.9 | 829 | 3% or 0.23 | 25 | 0.8 | 16 |
| *Fusarium graminearum* | 483 | 87 | growth | 0.9 | 197 | 20% or 0.015 | 40 | 0.8 | 9 |
| *Fusarium graminearum* | 483 | 87 | Disease severity | 0.9 | 197 | 20% or 6e-05 | 41 | 0.8 | 1 |
| *Eucaluptus grandis* | 1429 | 960 | Diameter | 0.99 | 823 | 10% or 0.01 | 143 | 0.4 | 6 |
| *Eucalyptus urophylla* | 1353 | 960 | height | 0.99 | 789 | 10% or 4.5e-05 | 138 | 0.4 | 1 |
| *Prunus avium* | 124/136 | 115 | BF | 0 | 123/135 | 20% or 0.020 / 20% or 0.004 | 25/27 | 0.4 / 0.4 | 3m/2m |
| *Prunus avium* | 124/136 | 115 | FF | 0 | 123/135 | 20% or 0.005 / 20% or 0.003 | 25/27 | 0.4 / 0.4 | 2m/1m |

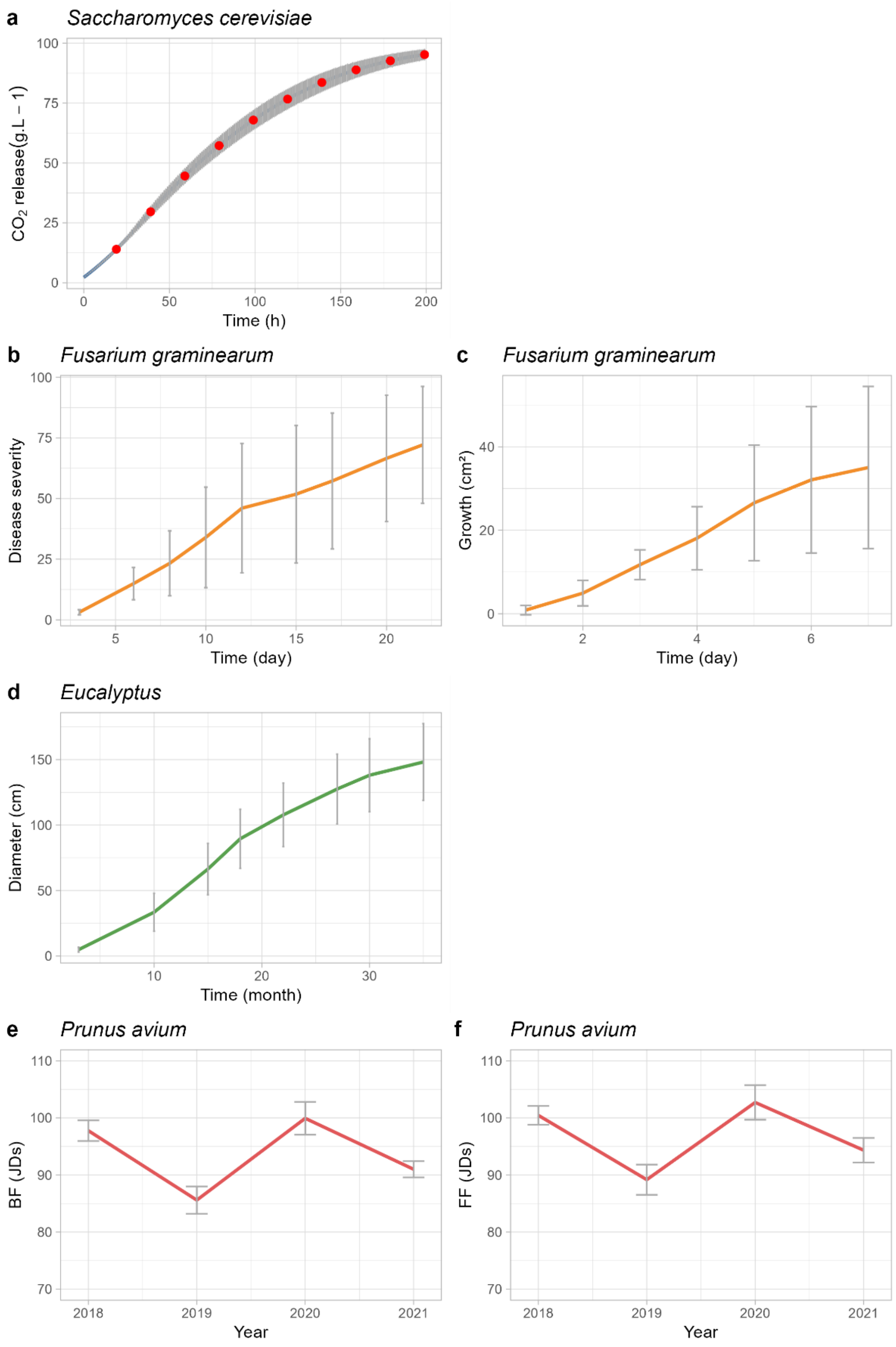


Figure S1: Mean of phenotypic value of the $CO_2$ release of *S. cerevisiae* (a), disease severity (b) and growth (c) of *F. graminearum*, tree diameter (d) of *Eucalyptus* hybrids, and beginning of flowering date (e) and full flowering date (f) of *P. avium* expressed in Julian Days (JDs) which are the number of days after the 1st of January of the considered year. The vertical lines represent the standard deviations at each time step. The blue, green, orange and red colors indicate the species *Saccharomyces cerevisiae*, *Eucalyptus (grandis × urophylla*, *Fusarium graminearum*, and *Prunus avium* respectively. For *S. cerevisiae* the ten-time steps considered in analysis are indicated by red points.

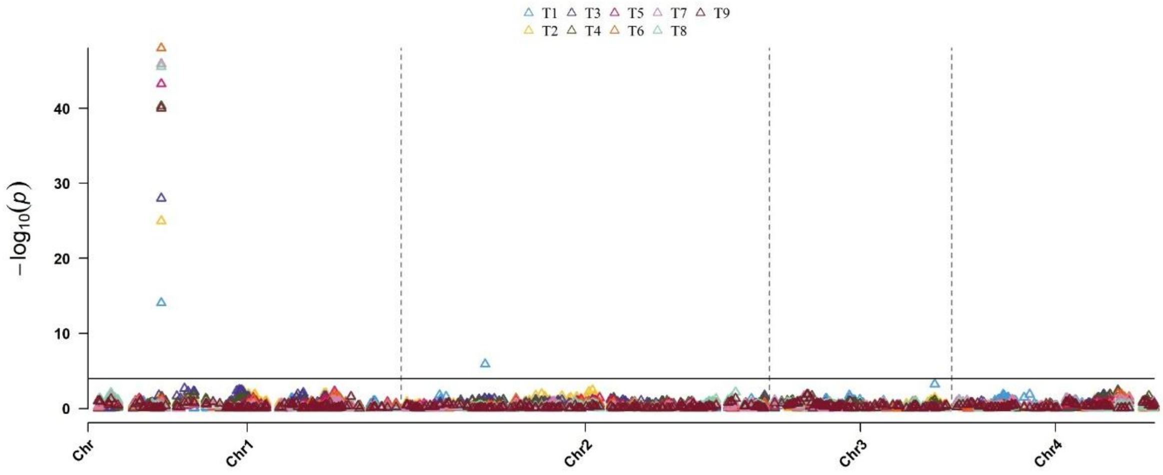


Figure S2: Multi-traits Manhattan plot of GWAS results for the 9 time steps of the disease severity time series of the *F. graminearum* data set. The results are issued from the BLINK model and each time step of the time series is represented by a specific color.

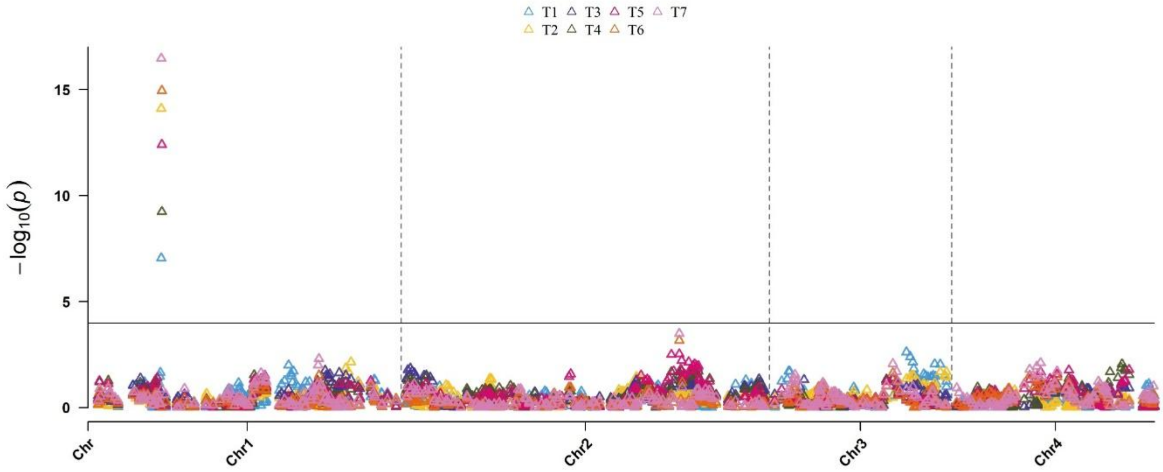


Figure S3: Multi-traits Manhattan plot of GWAS results for the 9 time steps of the growth area time series of the *F. graminearum* data set. The results are issued from the BLINK model and each time step of the time series is represented by a specific color.

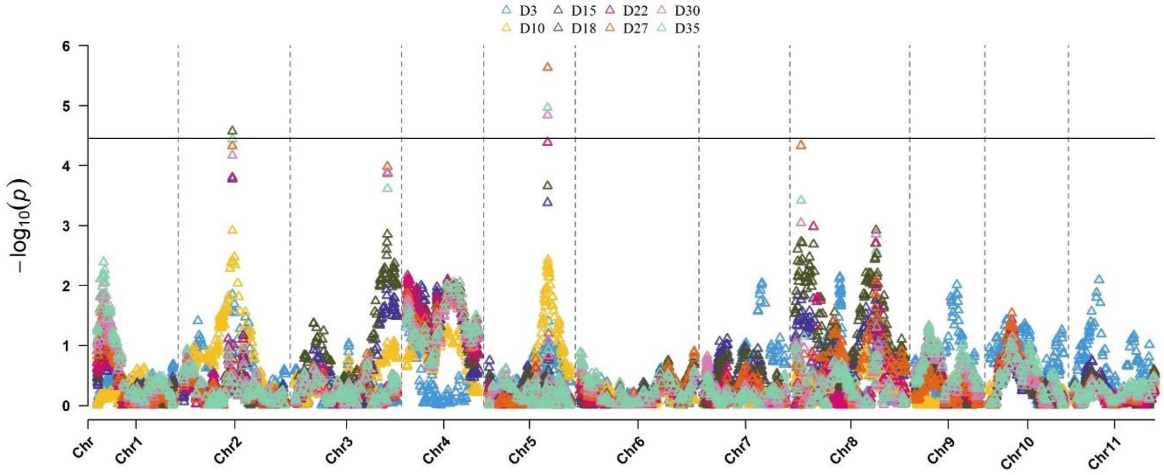


Figure S4: Multi-traits Manhattan plot of GWAS results for the 8 time steps of the diameter time series of the *E. grandis* data set. The results are issued from the BLINK model and each time step of the time series is represented by a specific color.

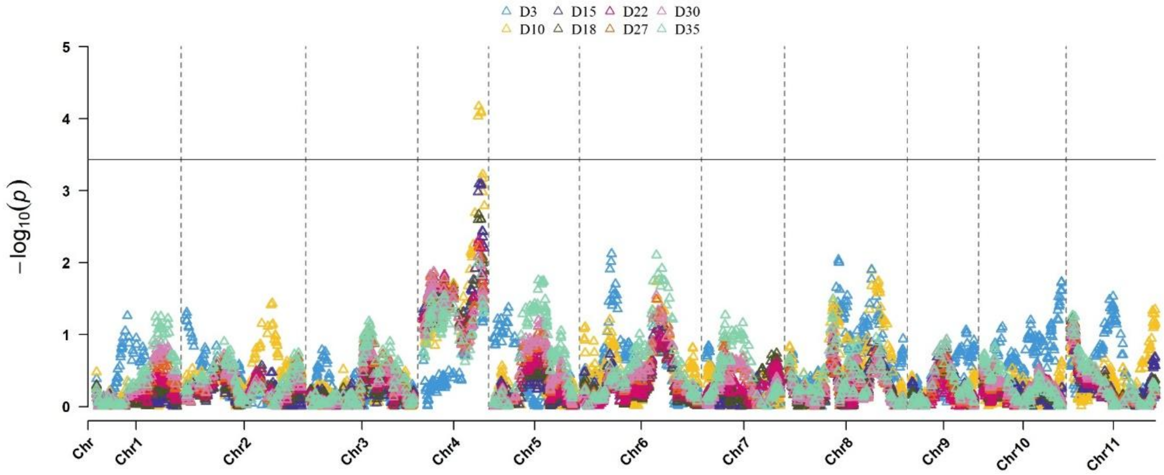


Figure S5: Multi-traits Manhattan plot of GWAS results for the 8 time steps of the diameter time series of the *E. urophylla* data set. The results are issued from the BLINK model and each time step of the time series is represented by a specific color.

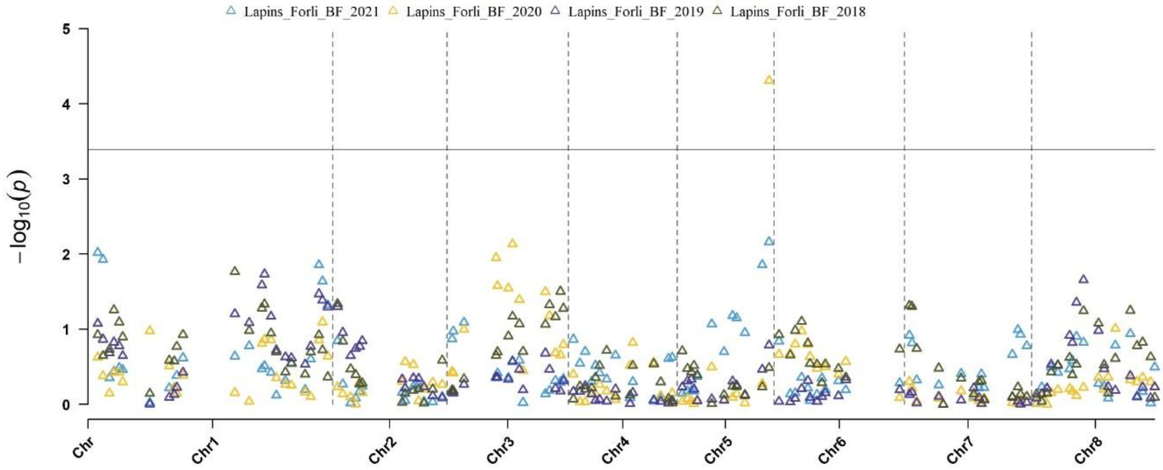


Figure S6: Multi-traits Manhattan plot of GWAS results for the 4 time steps of the beginning of flowering time series of the *P. avium* data set based on the Lapins genetic parental map. The results are issued from the BLINK model and each time step of the time series is represented by a specific color.

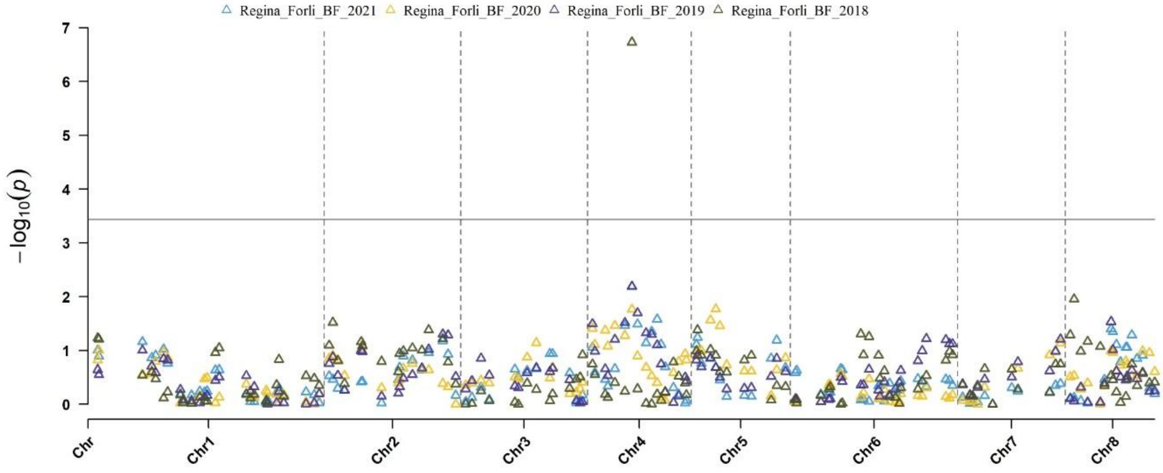


Figure S7: Multi-traits Manhattan plot of GWAS results for the 4 time steps of the beginning of flowering time series of the *P. avium* data set based on the Regina genetic parental map. The results are issued from the BLINK model and each time step of the time series is represented by a specific color.

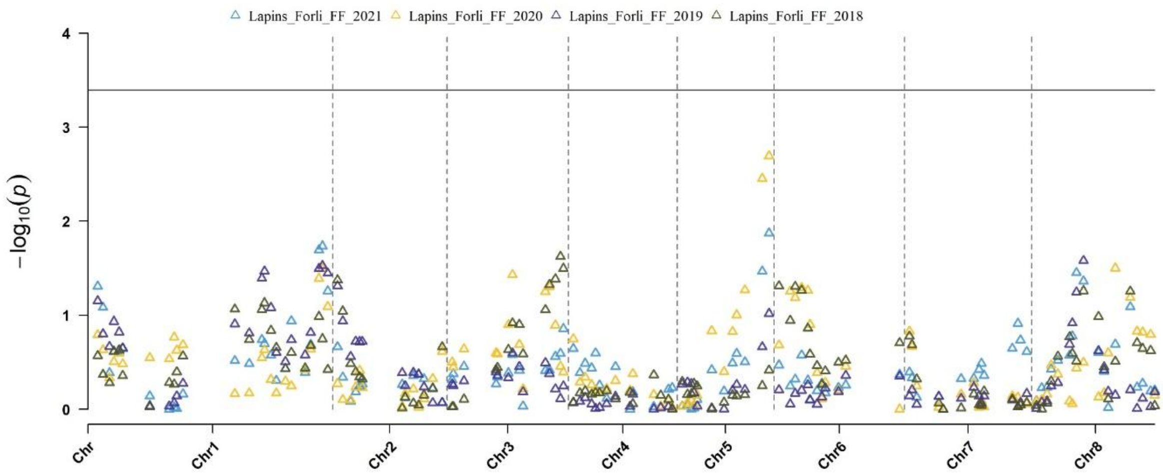


Figure S8: Multi-traits Manhattan plot of GWAS results for the 4 time steps of the full flowering time series of the *P. avium* data set based on the Lapins genetic parental map. The results are issued from the BLINK model and each time step of the time series is represented by a specific shape.

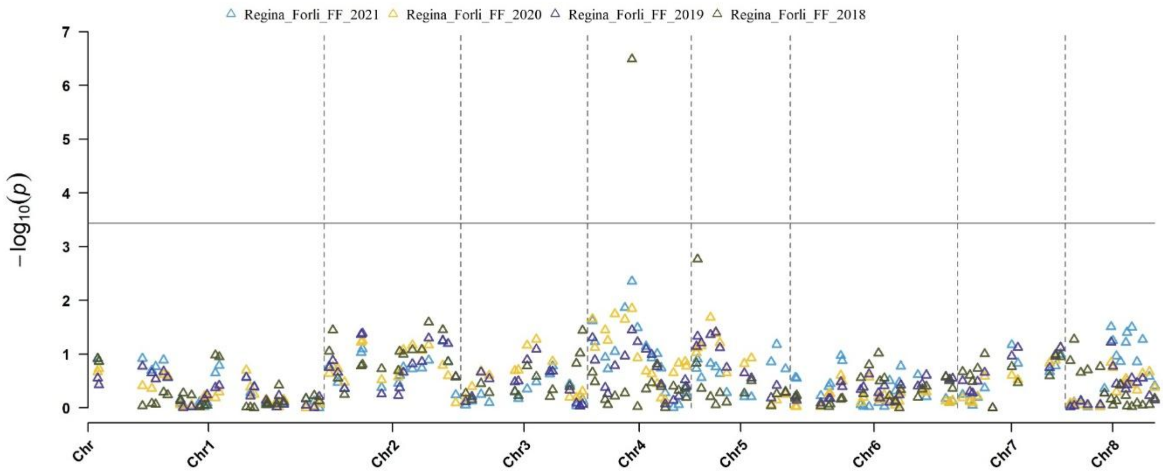


Figure S9: Multi-traits Manhattan plot of GWAS results for the 4 time steps of the full flowering time series of the *P. avium* data set based on the Regina genetic parental map. The results are issued from the BLINK model and each time step of the time series is represented by a specific shape.

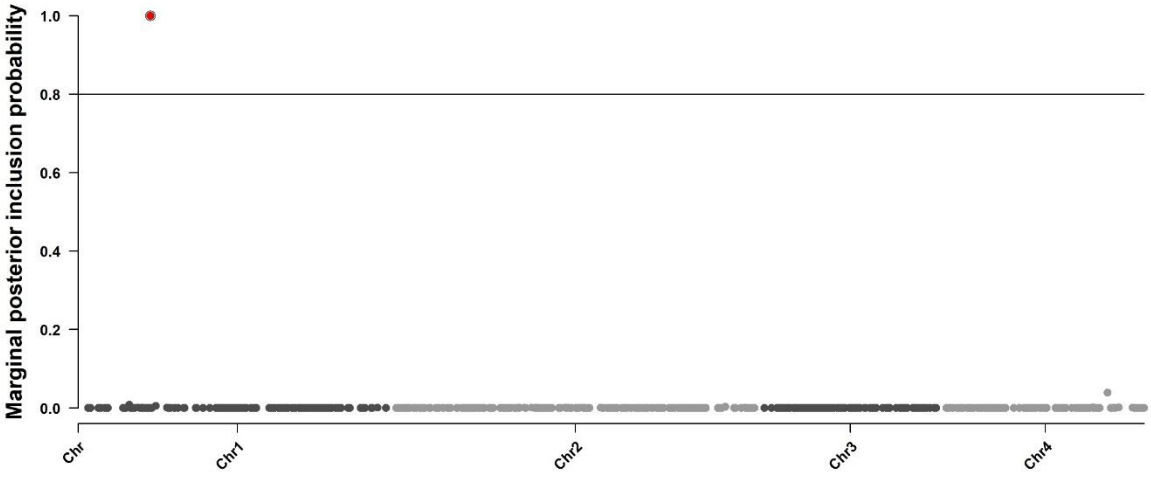


Figure S10: Marginal posterior inclusion probabilities of the *F. graminearum* genetic markers using the Bayesian Varying Coefficient Model. The probability is calculated according to the association between each marker and the variability of the disease severity time series. The line represent the selected threshold for marker selection and circles indicate the selected marker. Markers corresponding to genetic region also detected by the association mapping approach are filled in red.

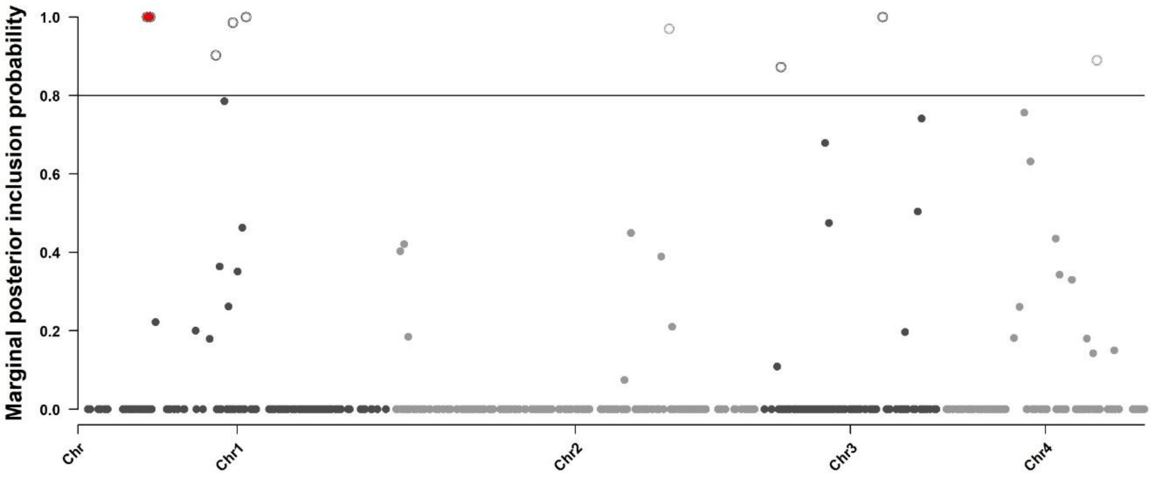


Figure S11: Marginal posterior inclusion probabilities of the *F. graminearum* genetic markers using the Bayesian Varying Coefficient Model. The probability is calculated according to the association between each marker and the variability of the growth area time series. The line represent the selected threshold for marker selection and circles indicate the selected marker. Markers corresponding to genetic region also detected by the association mapping approach are filled in red.

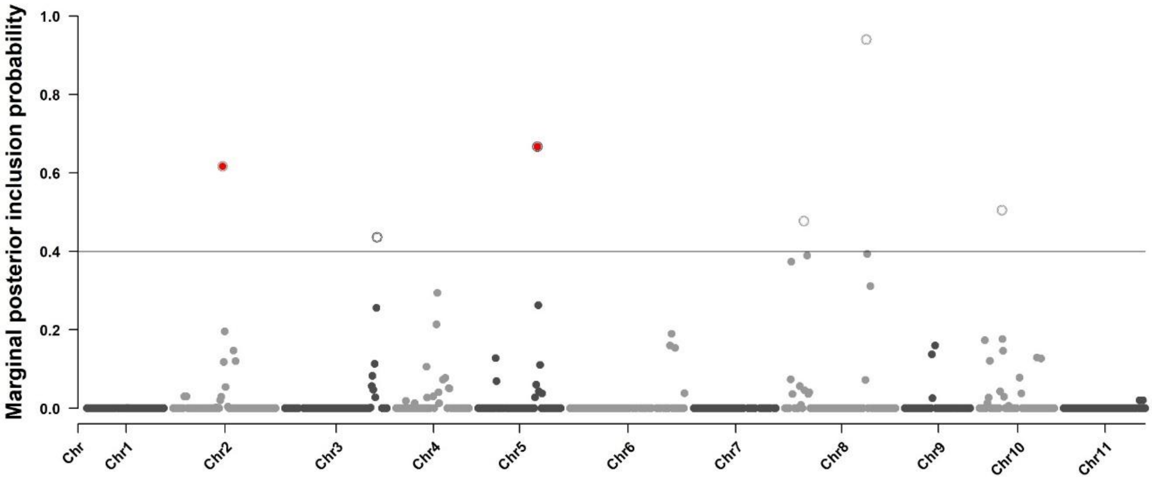


Figure S12: Marginal posterior inclusion probabilities of the *Eucalyptus* genetic markers, based on the *E. grandis* genetic parental map, using the Bayesian Varying Coefficient Model. The probability is calculated according to the association between each marker and the variability of the tree diameter time series. The line represents the selected threshold for marker selection and circles indicate the selected marker. Markers corresponding to genetic region also detected by the association mapping approach are filled in red.

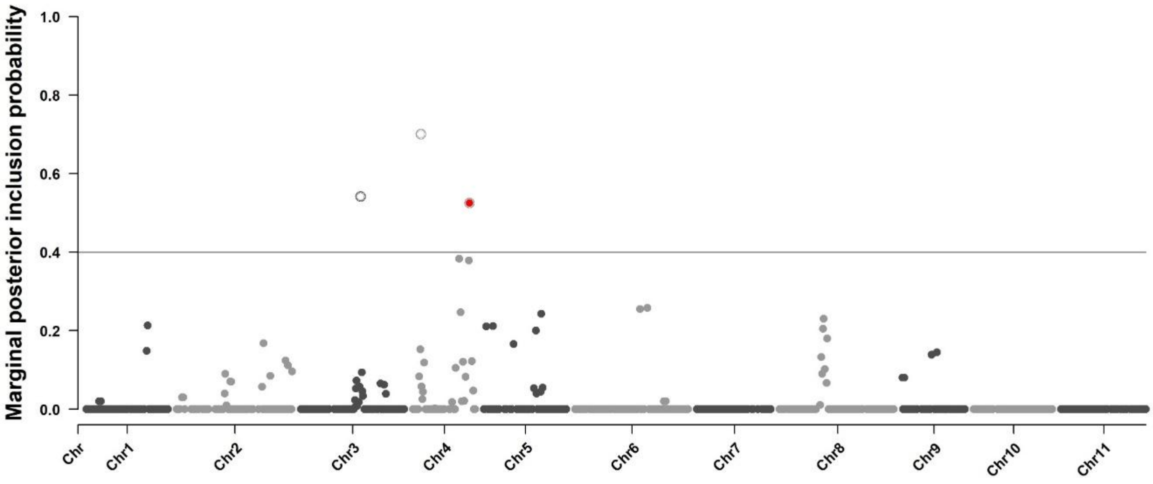


Figure S13: Marginal posterior inclusion probabilities of the *Eucalyptus* genetic markers, based on the *E. urophylla* genetic parental map, using the Bayesian Varying Coefficient Model. The probability is calculated according to the association between each marker and the variability of the tree diameter time series. The line represent the selected threshold for marker selection and circles indicate the selected marker. Markers corresponding to genetic region also detected by the association mapping approach are filled in red.

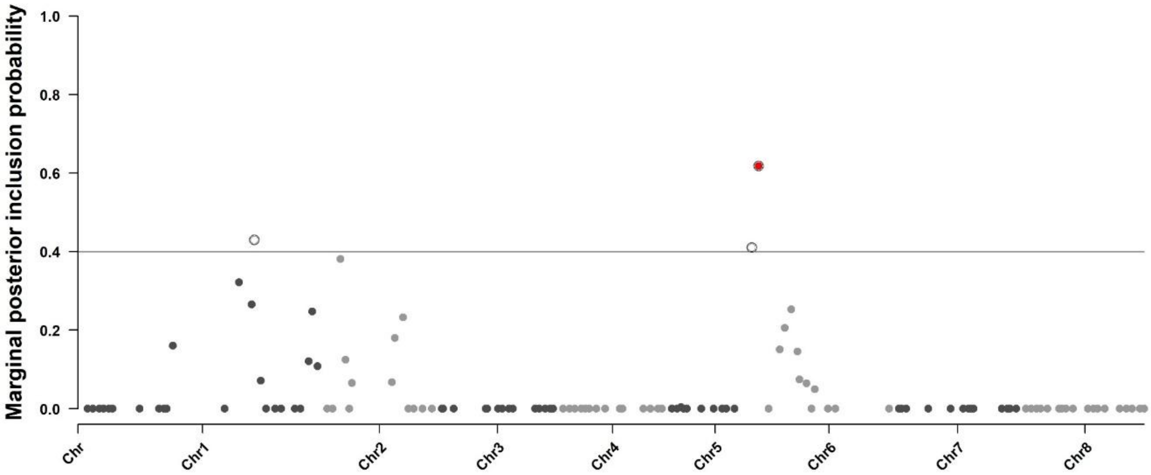


Figure S14: Marginal posterior inclusion probabilities of the *P. avium* genetic markers, based on the Lapins genetic parental map, using the Bayesian Varying Coefficient Model. The probability is calculated according to the association between each marker and the variability of the beginning of flowering time series. The line represent the selected threshold for marker selection and circles indicate the selected marker. Markers corresponding to genetic region also detected by the association mapping approach are filled in red.

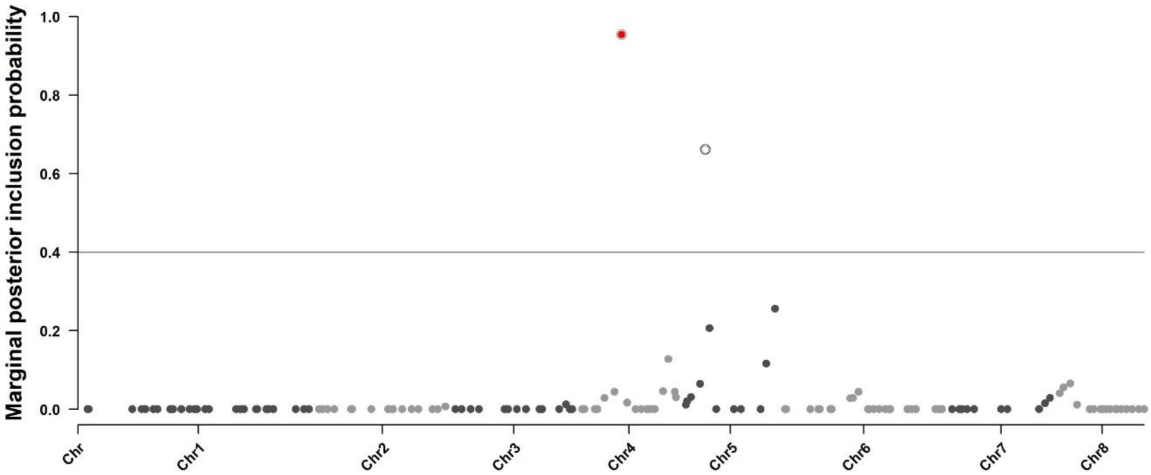


Figure S15: Marginal posterior inclusion probabilities of the *P. avium* genetic markers, based on the Regina genetic parental map, using the Bayesian Varying Coefficient Model. The probability is calculated according to the association between each marker and the variability of the beginning of flowering time series. The line represent the selected threshold for marker selection and circles indicate the selected marker. Markers corresponding to genetic region also detected by the association mapping approach are filled in red.

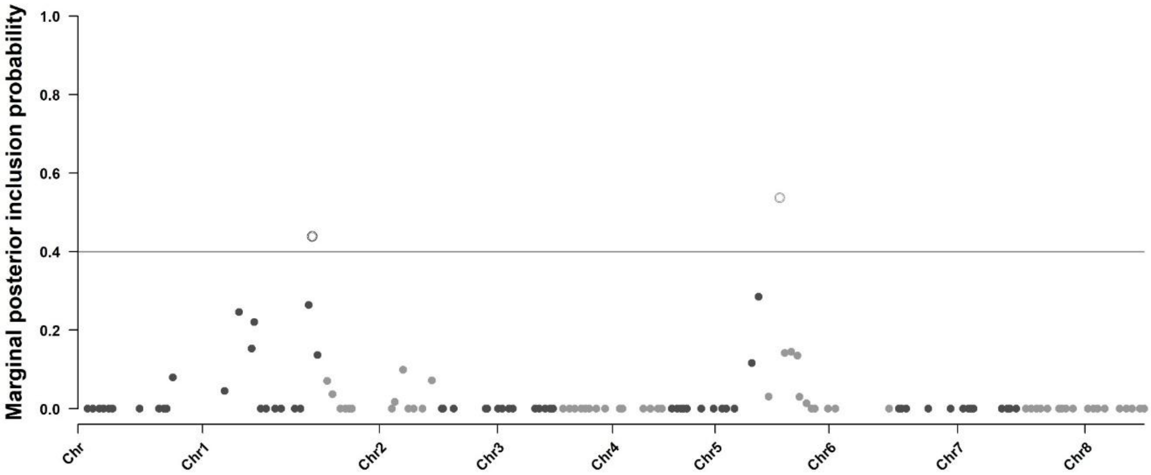


Figure S16: Marginal posterior inclusion probabilities of the *P. avium* genetic markers, based on the Lapins genetic parental map, using the Bayesian Varying Coefficient Model. The probability is calculated according to the association between each marker and the variability of the full flowering time series. The line represent the selected threshold for marker selection and circles indicate the selected marker. Markers corresponding to genetic region also detected by the association mapping approach are filled in red.

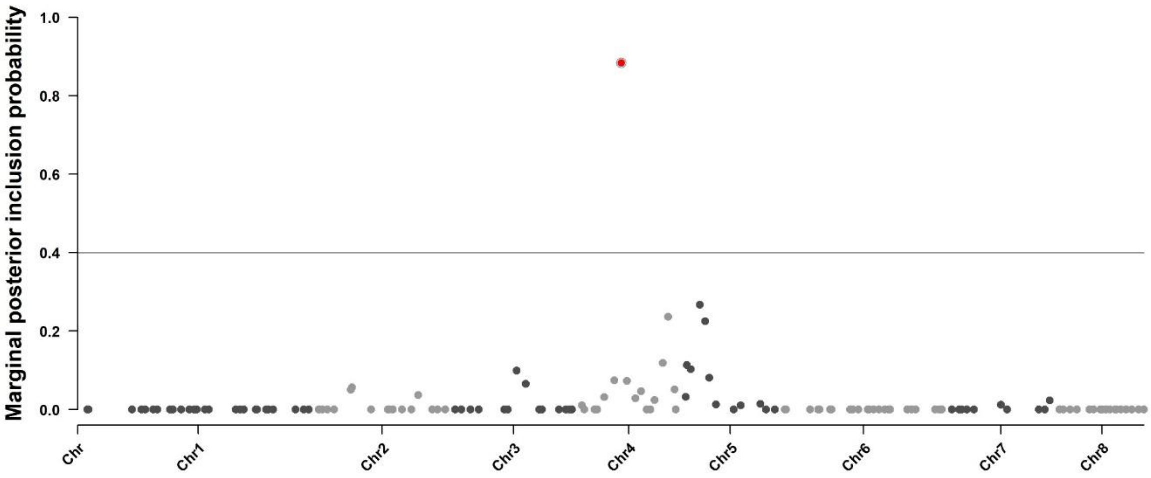


Figure S17: Marginal posterior inclusion probabilities of the *P. avium* genetic markers, based on the Regina genetic parental map, using the Bayesian Varying Coefficient Model. The probability is calculated according to the association between each marker and the variability of the full flowering time series. The line represent the selected threshold for marker selection and circles indicate the selected marker. Markers corresponding to genetic region also detected by the association mapping approach are filled in red.

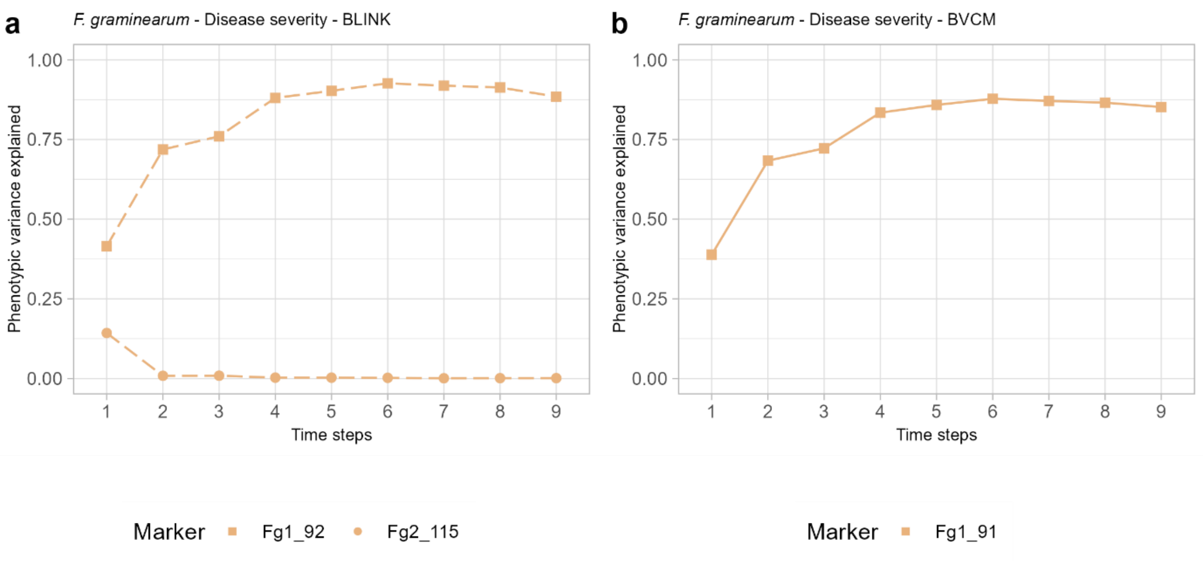


Figure S18: Phenotypic variance explained over time for the disease severity time series of *F. graminearum* by markers selected by the BLINK (dotted lines, a) and the BVCM (full lines, b) approaches. The markers commonly identified by BLIK and BVCM are indicated by light colored lines in b. Each marker is represented by a different point shape.

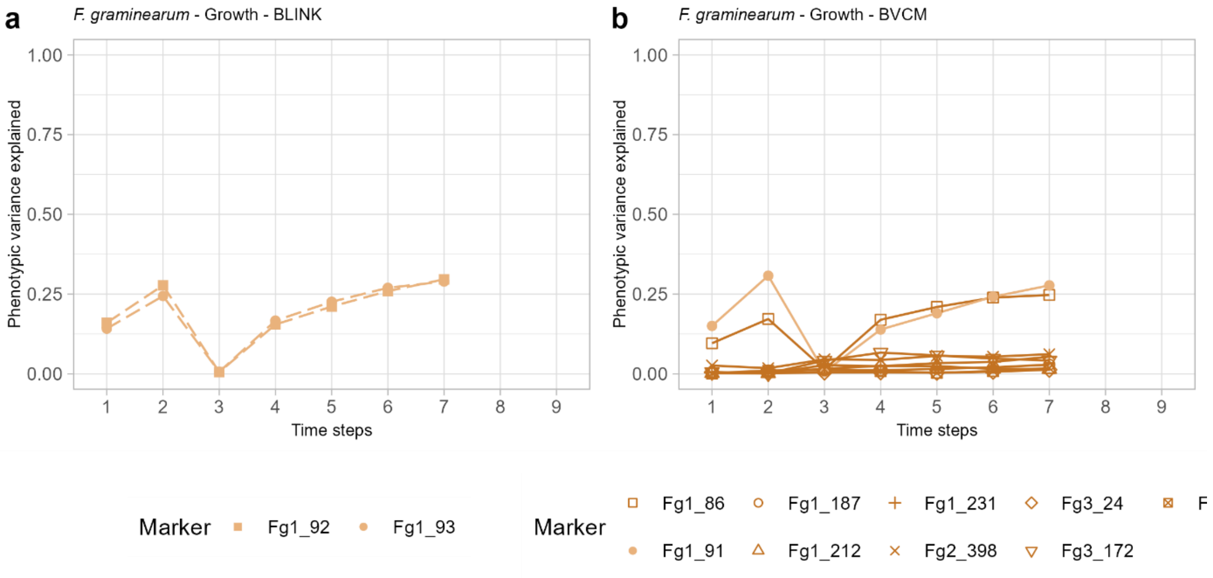


Figure S19: Phenotypic variance explained over time for the growth time series of *F. graminearum* by markers selected by the BLINK (dotted lines, a) and the BVCM (full lines, b) approaches. The markers commonly identified by BLIK and BVCM are indicated by light colored lines in b. Each marker is represented by a different point shape.

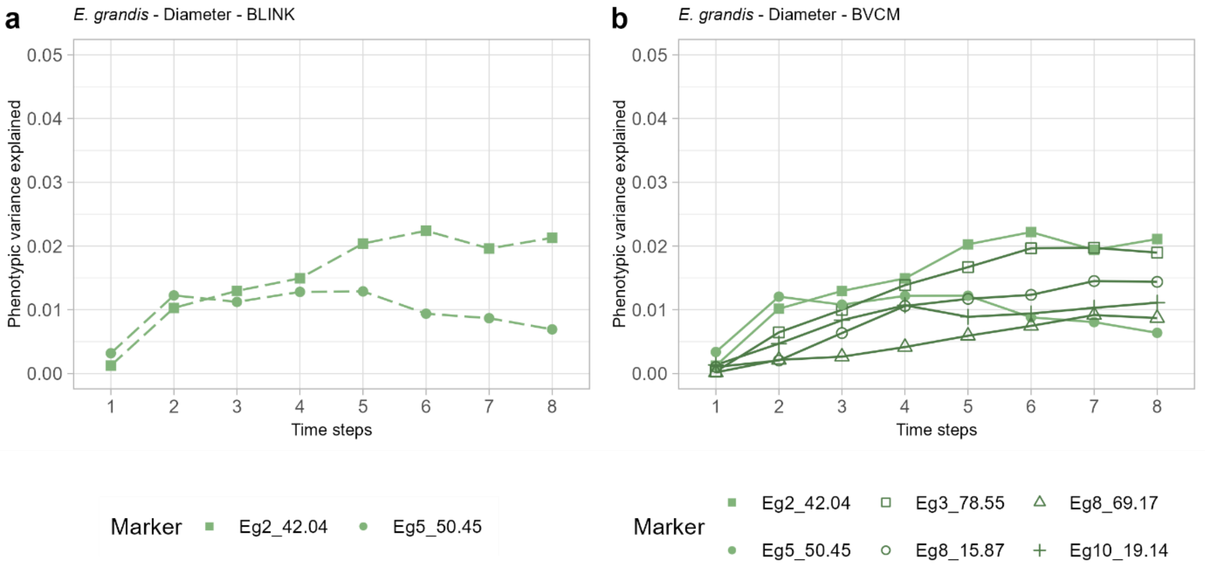


Figure S20: Phenotypic variance explained over time for the diameter time series of *E. grandis* by markers selected by the BLINK (dotted lines, a) and the BVCM (full lines, b) approaches. The markers commonly identified by BLIK and BVCM are indicated by light colored lines in b. Each marker is represented by a different point shape.

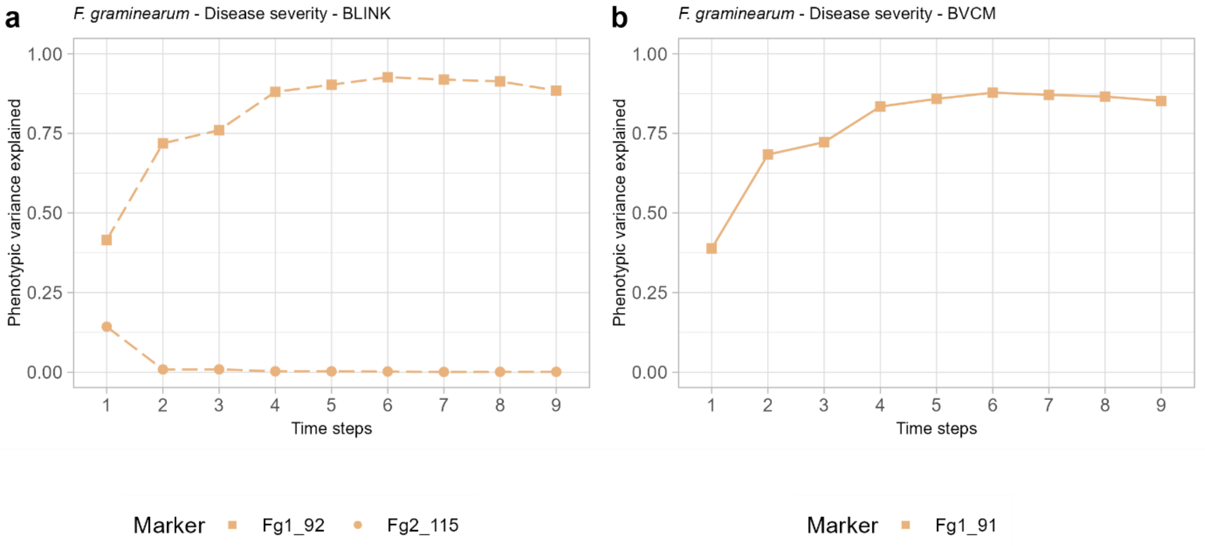


Figure S21: Phenotypic variance explained over time for the diameter time series of *E. urophylla* by markers selected by the BLINK (dotted lines, a) and the BVCM (full lines, b) approaches. The markers commonly identified by BLIK and BVCM are indicated by light colored lines in b. Each marker is represented by a different point shape.

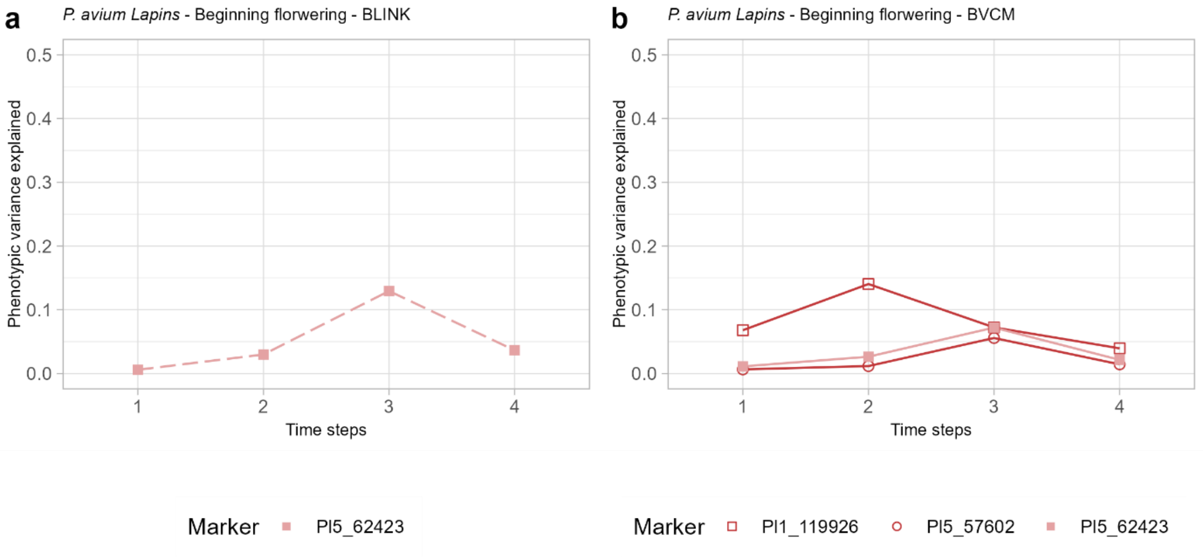


Figure S22: Phenotypic variance explained over time for the beginning of flowering time series of *P. avium* Lapins by markers selected by the BLINK (dotted lines, a) and the BVCM (full lines, b) approaches. The markers commonly identified by BLIK and BVCM are indicated by light colored lines in b. Each marker is represented by a different point shape.

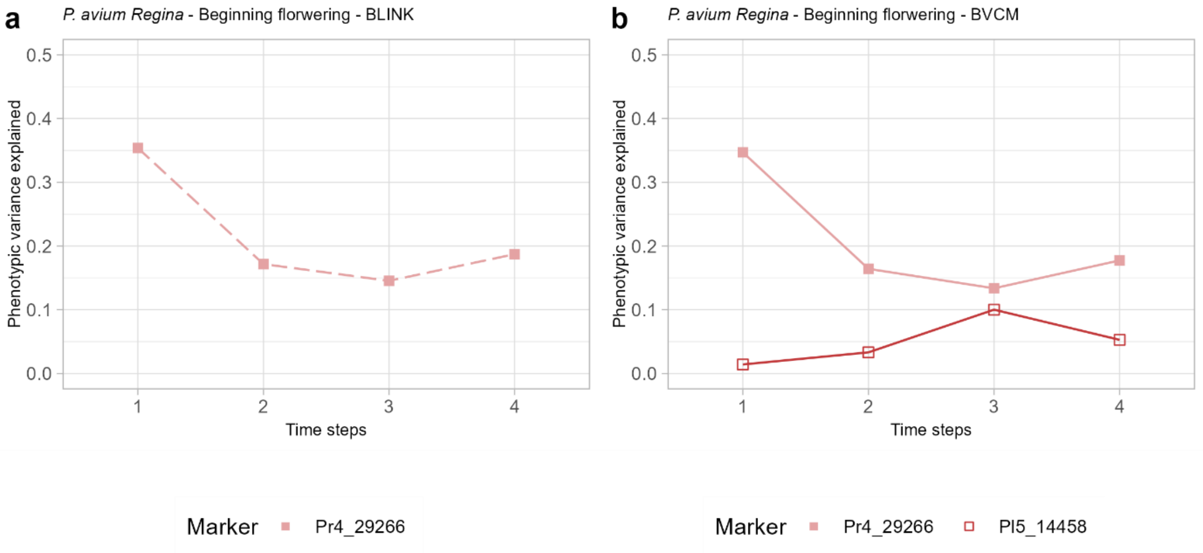


Figure S23: Phenotypic variance explained over time for the beginning of flowering time series of *P. avium* Regina by markers selected by the BLINK (dotted lines, a) and the BVCM (full lines, b) approaches. The markers commonly identified by BLIK and BVCM are indicated by light colored lines in b. Each marker is represented by a different point shape.

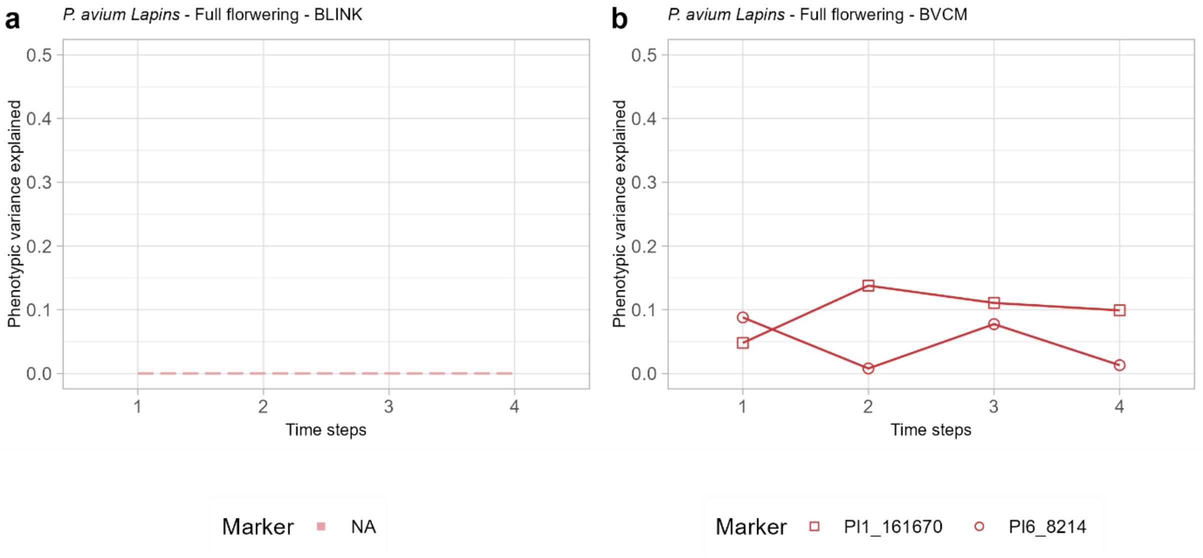


Figure S24: Phenotypic variance explained over time for the full flowering time series of *P. avium* Lapins by markers selected by the BLINK (dotted lines, a) and the BVCM (full lines, b) approaches. The markers commonly identified by BLIK and BVCM are indicated by light colored lines in b. Each marker is represented by a different point shape.

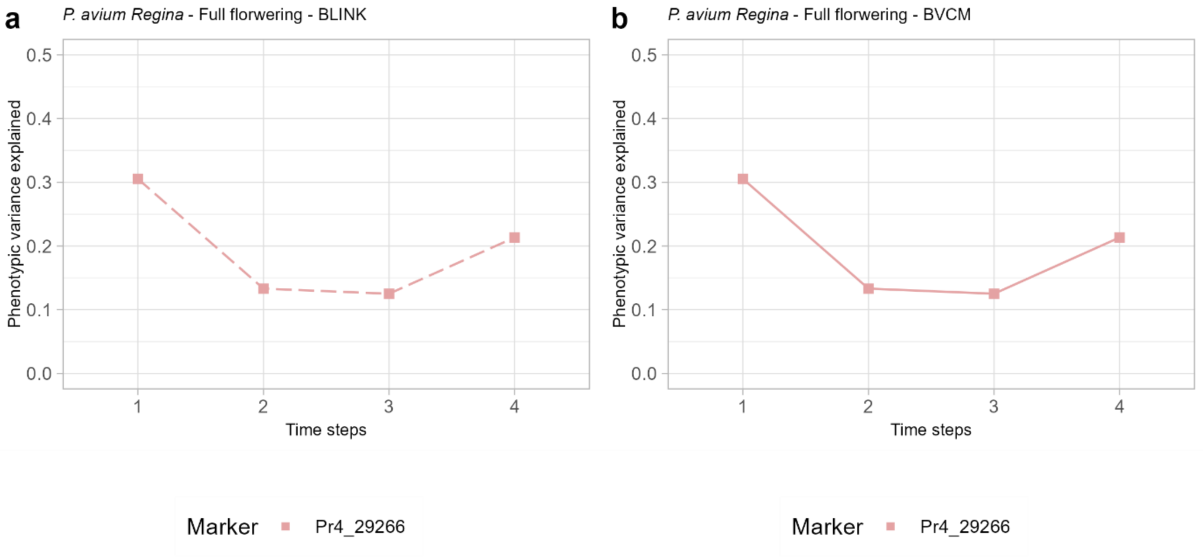


Figure S25: Phenotypic variance explained over time for the full flowering time series of *P. avium* Regina by markers selected by the BLINK (dotted lines, a) and the BVCM (full lines, b) approaches. The markers commonly identified by BLIK and BVCM are indicated by light colored lines in b. Each marker is represented by a different point shape.
